# Buckling of epithelium growing under spherical confinement

**DOI:** 10.1101/513119

**Authors:** Anastasiya Trushko, Ilaria Di Meglio, Aziza Merzouki, Carles Blanch-Mercader, Shada Abuhattum, Jochen Guck, Kevin Alessandri, Pierre Nassoy, Karsten Kruse, Bastien Chopard, Aurélien Roux

## Abstract

Many organs, such as the gut or the spine are formed through folding of an epithelium. This change in shape is usually attributed to tissue heterogeneities, for example, local apical contraction. In contrast, compressive stresses have been proposed to fold a homogeneous epithelium by buckling. While buckling is an appealing mechanism, demonstrating that it underlies folding requires to measure the stress field and the material properties of the tissue, which is currently inaccessible *in vivo*. Here we show that monolayers of identical cells proliferating on the inner surface of elastic spherical shells can spontaneously fold. By measuring the elastic deformation of the shell, we infer the forces acting within the monolayer and its elastic modulus. Using analytical and numerical theories linking forces to shape, we find that buckling quantitatively accounts for the shape changes of our monolayers. Our study shows that forces arising from epithelium growth in three-dimensional confinement are sufficient to drive folding by buckling.

## Introduction

Epithelium folding is essential for the formation of many organs, such as the gut during gastrulation and the central nervous system during neurulation (Davidson, 2012; Lecuit et al., 2011). There are several mechanisms that are typically considered to drive invaginations, in particular cellular flows and cell contractions (Davidson, 2012; Harris, 2017; Lecuit and Lenne, 2007). Specifically, apical constriction has been proposed to promote local invagination (Diaz-de-la-Loza et al., 2018). Moreover, islets of more contractile cells can form domains within a growing mesenchyme and lead to its folding (Hughes et al., 2018). All these mechanisms rely on a subpopulation of cells that have specific mechanical properties within the tissue (more contractile, more rigid, or more motile) and that locally generate mechanical stresses. Alternatively, the physical mechanism of buckling does not require heterogeneities or the generation of local stresses. Buckling is a bending instability occurring in elastic materials under compressive forces (Landau and Lifshitz, 1975). For example, a paper sheet lying on a table buckles when pushed inwards on opposite sides. Theoretical studies proposed that cell proliferation in confined geometries induce epithelium folding through buckling (Drasdo, 2000; Hannezo et al., 2014; Hocevar Brezavscek et al., 2012; Rauzi et al., 2015).

The shapes of many organs, for example, plant leaves (Liang and Mahadevan, 2009), villi in the mice gut (Shyer et al., 2013), and the drosophila wing (Tozluoğlu et al., 2019), are compatible with shapes resulting from buckling. However, it is required to measure the mechanical stresses in the epithelium and its mechanical properties in order to evaluate the specific contribution of buckling to epithelium folding. Several techniques are available for measuring mechanical stresses in single cells and tissues (Roca-Cusachs et al., 2017). A first category relies on local elastic deformation of the substrate balancing the internal forces of the tissue or cells, such as traction force microscopy (Mandal et al., 2014). Another category relies on mechanical probes embedded into the tissue, such as oil droplets (Mongera et al., 2018) or mechanosensitive lipid probes (Colom et al., 2018). Finally, these forces can be inferred from the dynamics of tissue deformations during morphogenesis, with some assumptions (Etournay et al., 2015; Guirao et al., 2015). Many of these techniques require knowledge of the tissue’s material properties, which are often impossible to measure directly.

## Results

### Cell encapsulation recapitulates epithelial monolayer folding under confinement

To directly test if mechanical stresses generated by proliferation can induce buckling of epithelia, we undertook an *in vitro* approach and studied the growth of a cell monolayer confined in a spherical shell. The spherical geometry presents several advantages over others: it is the one of early embryos, and it does not have boundaries, such that all cells experience the same three-dimensional environment. Specifically, we encapsulated MDCK-II cells in hollow alginate spheres, hereon referred to as capsules. To form them, we used a 3D-printed microfluidic device to generate three-layered droplets. Their outer layer consisted of alginate, which undergoes gelation in a 100mM CaCl_2_ solution (Fig 1a, Fig S1 and methods) (Alessandri et al., 2016; Alessandri et al., 2013). The inner surface of the capsules is coated with a 3-4 µm thick layer of Matrigel to which cells adhere (Fig 1g and Fig S1) (Alessandri et al., 2016) and on which they form epithelial monolayers (Fig 1b).

**Figure 1.**
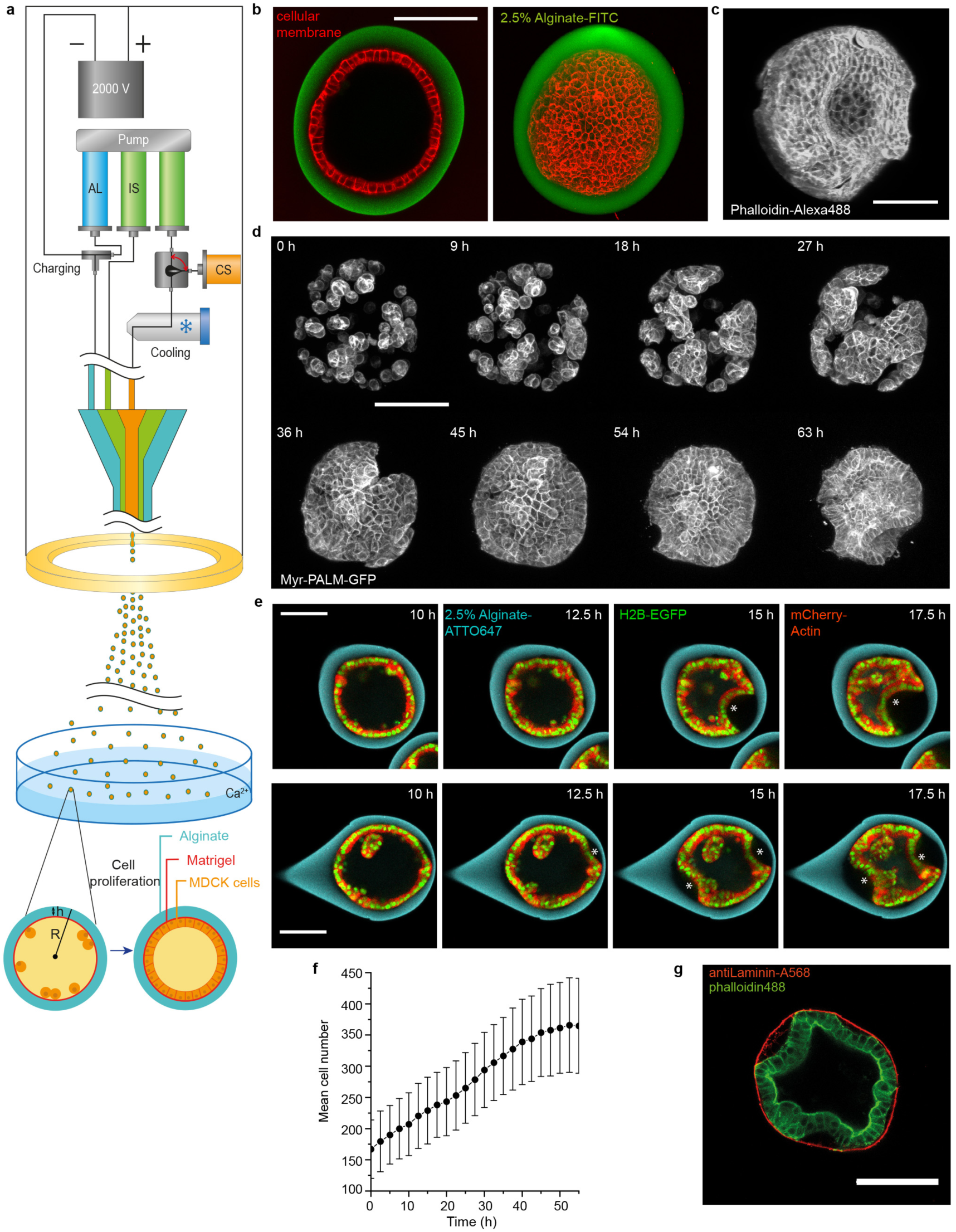
Characterization of epithelium growth and folding in a spherical capsule. (**a)** Schematic of experimental setup (see also Fig S1). (**b)** Left, confocal plane and right, maximum Z-projection of a fully formed MDCK spherical monolayer (Red, deep-red CellMask, membrane and green, FITC-alginate, capsule). (**c)** Maximum Z-projection of a fixed and folded MDCK monolayer stained with Phalloidin-Alexa488 (actin), respectively. (**d**) Maximum projections of confocal planes of MDCK-Myr-PALM-GFP cells forming spherical epithelial monolayer that folds after 45h. (**e**) Confocal equatorial planes of one fold (top) and two folds (bottom) of epithelium. Asterisks indicate folds. (**f**) Mean cell number per capsule over time; 3 experiments, N=53 capsules; error bars are SDs. (**g)** A confocal equatorial image of a folded MDCK II monolayer after fixed and immunostaining with antiLaminin-A568 (red, Matrigel) and phallodin-Alexa488 (green, actin). Scale bars, 100 µm.

Encapsulated cells were imaged using 3D time-lapse confocal microscopy. Time zero corresponds to the start of imaging, 24h after capsule formation (Methods), unless mentioned otherwise. Initially, MDCK-II cells were sparsely distributed on the capsule’s inner surface. Through proliferation, cells first formed clusters, which then merged into a monolayer (Fig 1d). Monolayers reached confluency at 8.8±0.8 hours (mean±SEM, as in the rest of the text, unless noted, N=54). Monolayers folded after 14.5±0.8 hours (N=54) in approx. 80-90% of the capsules. In this process, a portion of the monolayer detached from the alginate shell and progressively bent inwards (Fig 1c, d, e, SI Movies 1). Proliferation was unaffected by confluency or folding, as the cell number increased linearly with a rate of 3.6±0.1 cells per hour (N=54) for 55 hours (Fig 1f), consistent with the established growth dynamics of MDCK cells (Soderberg et al., 1983). From these observations, we concluded that cell monolayers confined in a spherical shell can spontaneously fold.

### Epithelial monolayer folding is due to buckling

Before investigating in detail whether epithelium folding in our experiments was due to buckling, we checked whether other processes contributed to folding. First, monolayers of MDCK-II cells on micro-patterned substrates have been reported to form osmotically swelling domes by ionic pumping through the cell monolayer (Latorre et al., 2018). To sustain the osmotic pressure difference that grow the domes, the substrate needs to be impermeable to these ions, and the monolayer must have no holes. In our experiments, the monolayer remained sealed during folding as evidenced by negligible fluorescein loss from the monolayer lumen (Fig S2). The exchange of volume between the lumen of the monolayer and its exterior during folding can be accounted for by the flux of water through the monolayer (SI Section 1). In contrast to the experiments in (Latorre et al., 2018), where PDMS was used as a substrate, alginate shells cannot maintain significant osmotic pressure differences: alginate is highly permeable to small molecules, as we showed by fast fluorescein diffusion from the capsule exterior to its interior (Fig S2).

Second, in some morphogenetic processes, active cell flows induce sufficient stresses to fold epithelia (Etournay et al., 2015; Munster et al., 2019). In our assay, however, no large-scale collective cellular flows were observed (Fig 2b). Altogether, we concluded that none of these mechanisms explained monolayer folding in our assay and we further investigated the possibility of epithelium buckling due to stresses by cell proliferation under confinement.

**Figure 2.**
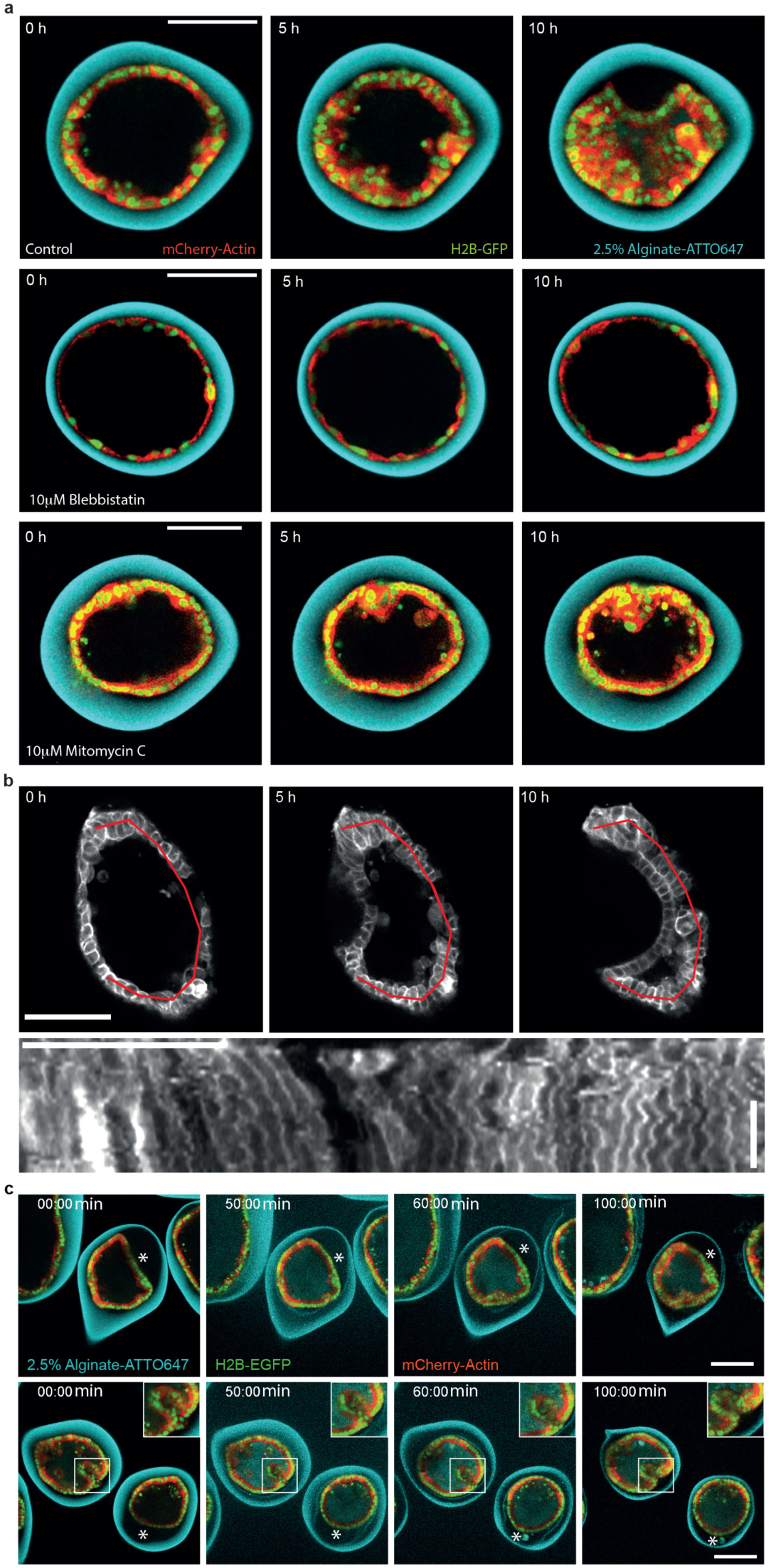
Monolayer folding is due to buckling. (**a**) Confocal equatorial planes of MDCKII monolayers. Top, non-treated control; middle, application of 10 μM Mitomycin-C; bottom, application of 10 μM blebbistatin. T=0 corresponds to cell confluency (**b**) Top; Confocal equatorial planes of Myr-PALM-GFP-MDCK monolayer during folding. T=0 corresponds to the beginning of folding. Bottom; Kymograph along the red line shown above 10h post-folding. Time is towards the bottom (see arrow on the right). Time scale bar, 5h. (**c**) Confocal equatorial planes of MDCKII monolayer relaxation due to capsule dissolution with alginate lyase. Asterisks highlight relaxation. Scale bars, 100 µm.

Finally, acto-myosin contractility can lead to folding, for example, through apical constriction or in presence of differences between basal and lateral tension(Sui et al., 2018). Since the apical side of our encapsulated monolayers faced the interior of capsules (Fig S3), apical constriction would rather oppose than promote folding in our system (Diaz-de-la-Loza et al., 2018; Krajnc and Ziherl, 2015; Lecuit and Lenne, 2007; Storgel et al., 2016). We further investigated the role of acto-myosin contractility for folding by treating capsules with 10 μM blebbistatin to inhibit Myosin II. In that case, cells did not detach from the alginate shell, and no folds emerged (Fig 2a, Movie 2). This result was fully consistent with our recent work showing that contractility of epithelial cells was required to detach the cell monolayer encapsulated in alginate tubes (Maechler et al., 2019). In these alginate tubes, the epithelium detached but did not fold, and blebbistatin inhibited detachment. In those tubes, cell monolayers did not fold because cells were not confined, as they can grow along the tube axis almost indefinitely. We however wanted to test further if acto-myosin constriction was essential to folding in the spherical capsules after detachment.

To bypass the effect of confinement, we blocked cell proliferation. To this end, we treated encapsulated epithelia with 10 μM Mitomycin C to block their proliferation at the time they reached confluency. After ten hours of Mitomycin treatment no folds were observed, whereas epithelia were folded in control capsules (Fig 2a, Movie 3). On longer time, cell monolayers detached, teared and reorganized but never folded (Fig S4). These results showed that when proliferation is blocked, and contractility is not affected, monolayer detachment is observed, but no folding, supporting our previous findings that cell contractility is involved in detachment and not in folding (Maechler et al., 2019). It also suggests that confinement is essential to folding.

To further check that confinement was essential, we dissolved capsules using alginate lyase immediately after monolayers reached confluency. In this case either, we could not observe monolayer folding (Fig 2c, Movie 4). We concluded that confinement is essential for monolayer folding. Interestingly, if capsules were dissolved after folding, epithelia just detached or partially folded relaxed to a round shape, while fully folded epithelium only partially relaxed (Fig 2c). On the longer overnight time scale, fully formed folds almost fully relaxed. This suggested that epithelia were under compressive stresses during folding and supported the buckling hypothesis.

### Measurements of compressive stresses due to monolayer proliferation and folding

Since alginate is elastic, stresses caused by cell proliferation can be inferred from capsule deformations: the alginate shell undergoes expansion and thinning during proliferation (Fig 3a). Capsules without cells kept a constant wall thickness (Fig S5). To infer the effective pressure corresponding to these compressive stresses, we measured the average capsule radius and wall thickness in confocal images (Fig 3b-d), and their elasticity modulus (Young’s modulus) by atomic force microscopy (AFM) (Fig 3e and methods), for four different alginate concentrations (1, 1.5, 2, and 2.5%, see methods, Fig S5). For all alginate concentrations, the pressure increased during approximately 55h, reaching a maximal value of 300-400 Pa (Fig 3f). The proliferation rate being constant during the 40-50 first hours, we concluded that pressures below 300 Pa did not significantly affect the proliferation rate.

**Figure 3.**
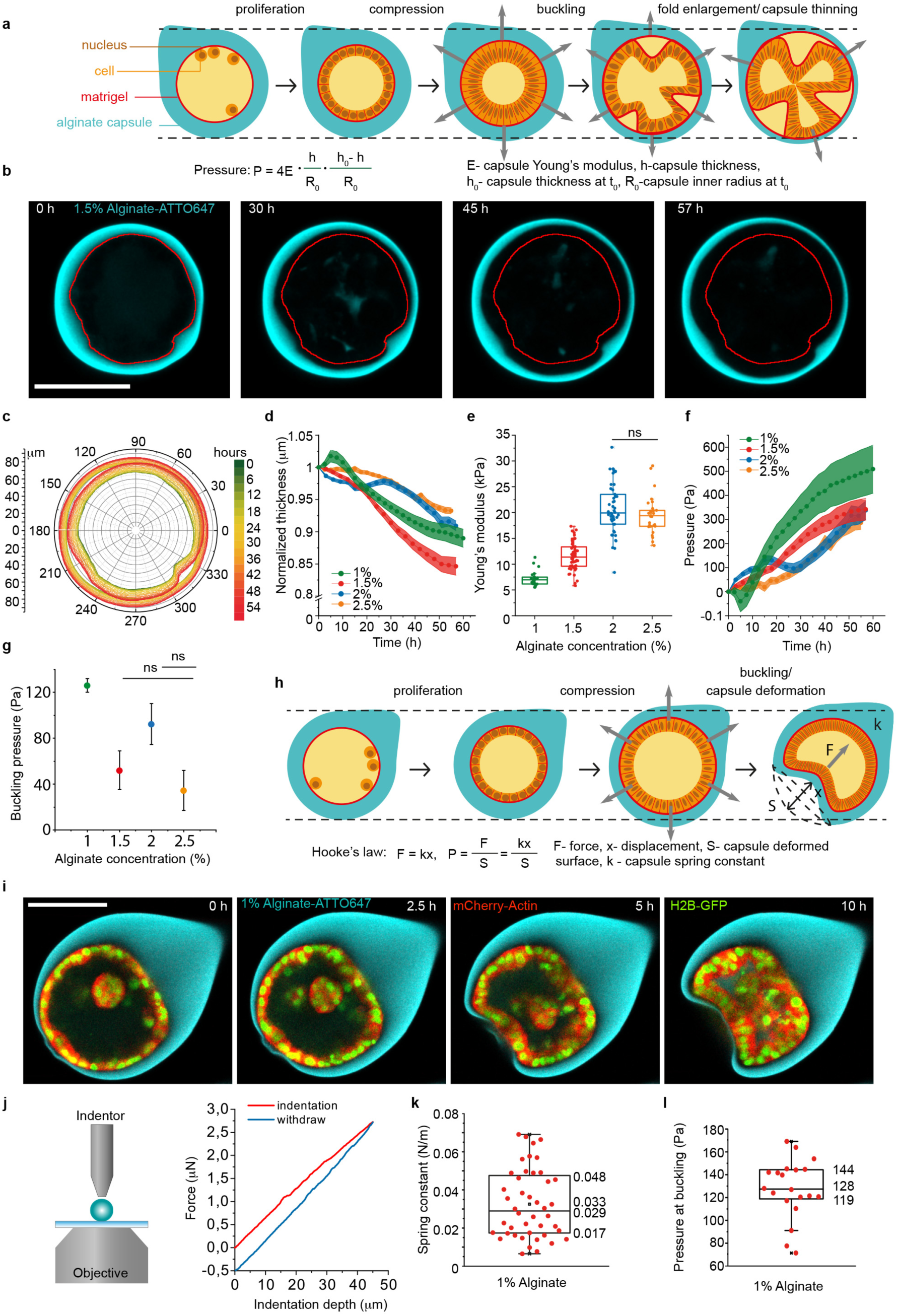
Measurements of compressive stresses due to monolayer proliferation and folding. (**a**) Schematic of capsule thinning during epithelium proliferation. (**b**) Confocal equatorial plane of thinning alginate capsule. Red contours correspond to the capsule inner perimeter at T=0h. (**c**) Superimposed contours of inner and outer boundaries corresponding to different time points. (**d**) Normalized mean capsule thickness as a function of time for different alginate concentrations. (**e**) Young’s modulus (kPa) as a function of the alginate concentration measured by AFM. Respective Young moduli are: 1%, 7.1±0.3 kPa (N=25), 1.5%, 11.5±0.4 kPa (N=52), 2%, 20.7±0.7 kPa (N=46) and 2.5%, 19.5±0.7 kPa (N=29). Difference between 2% and 2.5% alginate is not statistically significant (ns) with two-tailed P value 0.7042. (**f)** Evolution of pressure (Pa) within capsules over time during epithelium proliferation and for different alginate concentrations (see methods). (**g)** Mean buckling pressure (Pa) for different alginate concentration. For d, f, and g: 1% alginate, n=22; for 1.5%, n=35; for 2%, n=25; for 2.5%, n=53; error bars are SEM. Difference between 1.5% and 2% alginate, and between 2.5% and 1.5% are not statistically significant (ns) with two-tailed P values 0.1119 and 0.4909, respectively. (**h**) Schematic of capsule invagination following epithelium folding. (**i**) Confocal equatorial planes showing capsule invagination for 1% alginate capsule. (**j**) Left, schematic of an indentation experiment with the FemtoTools indenter (see also Fig S5). Right, a representative plot of force with indentation depth. (**k**) Box plot of spring constant for 1% alginate. (**l**) Box plot of pressure at buckling (Pa) for 1% alginate capsule calculated from capsule deformation (see methods). Scale bars, 100 µm.

The pressure at folding measures stresses required for folding, and was between 50 and 100 Pa for 1.5, 2 and 2.5% alginate concentration (Fig 3g). Thus, it did not significantly depend on the capsule stiffness. For 1% however, the monolayer did not detach from the capsule shell upon folding in approximately 65% of the cases. Rather, the shell was pulled in by the folding monolayer (Fig 3h-i, Movie 5) confirming that osmotic pressure was not at the origin of folding. We estimated the force exerted by cells onto 1% alginate shells by using Hooke’s law for the deformation of the capsule (Fig 3h). The effective spring constant (0.03±0.003 N/m, N=44) was measured using an FT-S100 indenter (FemtoTools, Buchs, Switzerland, see methods, Fig S5 and Fig 3j-k). This value results in an average maximal deflection force of 1.8±0.2 μN (N=20), about a hundred time larger than the maximal force exerted by a single cell (Mandal et al., 2014). Dividing this value by the invagination area, we obtained a pressure of 128±6 Pa (N=20) (Methods, Fig 3l), which is of the same order as for other alginate concentrations (Fig 3g).

### Continuum theory of the buckling transition supports buckling

We theoretically determined the compressive stresses within the monolayer corresponding to the pressure at folding. To this end, we used a continuum description without cellular details. Cell monolayers are described as a circular elastic ring in two dimensions, reproducing the geometry in an equatorial confocal plane (Fig 4a-b and SI section 2). Confinement is accounted for by restricting the cellular ring to a circular domain of radius R. If not confined, the ring is circular and has a radius *r*. If confined and *r>R*, then the ring is deformed, namely, compressed or bent. Two elastic parameters characterize its resistance to these deformations: the bending rigidity *𝒦* and the compressional rigidity λ (SI section 2). For *r>R*, confinement is achieved by a harmonic force of spring constant *k*, which represents the elasticity of the capsule. The ring shape is determined by minimization of the total energy, which is a combination of the bending, compression, and confinement energies (SI section 2).

**Figure 4.**
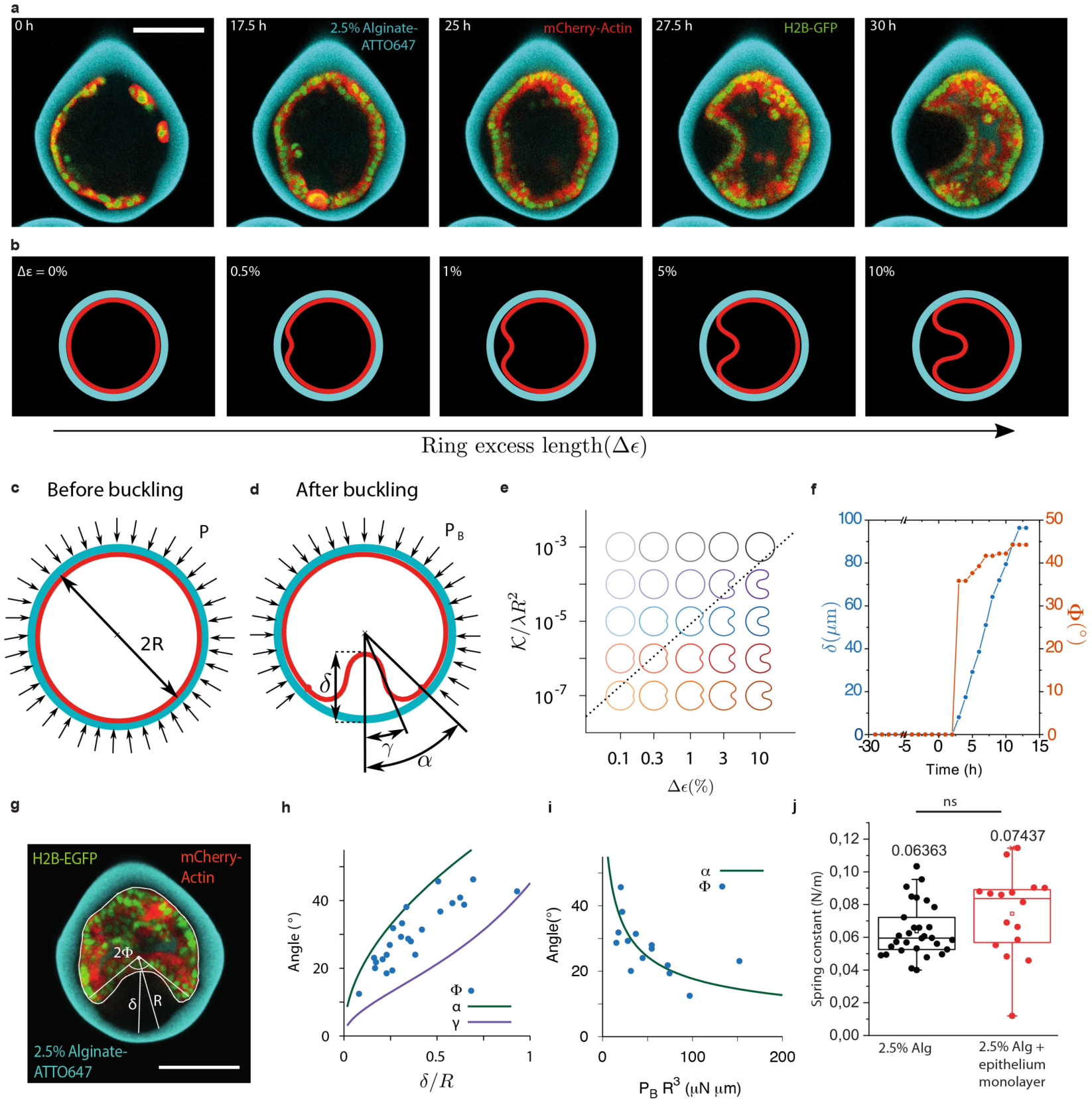
Continuum theory of the buckling transition and comparison to experimental data. (**a)** Confocal equatorial planes of epithelial monolayer bending. (**b**) Equilibrium shapes of a buckled elastic ring (red) under circular confinement (cyan) as a function of ring excess strain (Δε), calculated from continuum theory (see text and SI). (**c**) Schematic of a compressed elastic ring (red) under the pressure P of the confinement ring (cyan). (**d)** Schematic of a buckled elastic ring (red) under the pressure P_B_ of the confinement ring (cyan). (**e)** Equilibrium shapes as a function of *𝒦* /λR^2^ and the excess strain Δε. The dashed line stands for the threshold given by eq. S25. List of parameters: *𝒦*=10^−2^, k=10^5^, R=1 and λ varies from 10 to 10^5^. (**f)** Experimental values of δ and Φ as a function of time. 0 time point corresponds to the monolayer confluence. (**g**) Experimental measurements of δ, Φ, R depicted on a confocal equatorial scan of a capsule with a buckled monolayer. (**h)** Blue dots, experimental values of (Φ;δ). Solid lines; theoretical relations between δ and α (green) and δ and γ (purple) for the compressional rigidity 10^8^. (**i**) Blue dots, experimental values (Φ,P_B_ R^3^) from panel **g**. Only dots where δ was smaller than R were kept for the fit. Solid green line fits to the theoretical relation between α and P_B_R^3^ giving *𝒦*=0.5 μN.μm. (**j**) Spring constant values of 2.5% alginate capsules with and without a cell monolayer (see Fig S5). The difference is not statistically significant with two-tailed P value 0.5809.

In our theory, we capture monolayer size by the excess strain Δε=ΔL/2π*R*, where ΔL=2π(*r-R*) is the excess length (Fig 4b). This system exhibits a first order buckling transition controlled by the excess strain Δε (Chan and McMinn, 1966; Lo et al., 1962) (Fig S6 and SI, Section 3). If the rigidity of the capsule is much higher than the one of the epithelium (λ/*k*R^2^<1), the threshold excess strain at the transition can be deduced by comparing the energies in two limiting cases: a compressed unbuckled ring (Fig 4c) and an uncompressed buckled ring (Fig 4d). In the first case (Fig 4c), stationary rings accommodate Δε through compression, resulting in circular shapes. The local strain scales as ∼ Δε and the compressed ring length as ∼ R, resulting in a compressive energy per unit length ΔE ∼ λ*R*Δε^2^ (SI Section 3). In the second case (Fig 4d), rings accommodate Δε through bending deformations. We estimate the energy of a buckled ring by splitting it into two parts: a folded segment and an undeformed ring segment (Fig 4d). The fold is characterized by its depth δ and its opening angle α (Fig 4d, Fig S6). Minimizing the energy for this shape while keeping the total length constant, yields α ∼ Δε^1/3^ and δ ∼ *R*Δε^2/3^ (SI Section 3). The fold average curvature scales as ∼ δ/(Rα)^2^ and the folded segment length scales as ∼ *R*α, resulting in a bending energy per unit length of the buckled ring ΔE ∼ *𝒦*Δε^1/3^/*R* (SI, Section 3). The buckling transition occurs when the energies of both the constraint ring and the buckled ring are comparable, which happens for a threshold excess strain Δε_c_ ∼ (*𝒦/*λR^2^)^3/5^. This threshold depends on both, the tissue material properties and the confinement geometry (Fig 4e and SI Section 3).

From this, the pressure at buckling can be determined. Below the threshold (Δε<Δε_c_), the ring compression is stabilized by a uniform pressure P exerted by the confinement (Fig 4c). The energy associated with this pressure is ∼P*R*^2^Δε and equals the ring compression energy ΔE∼λ*R*Δε^2^, yielding P∼λΔε/*R* (Fig S6 and SI Section 4). Hence, the pressure at the buckling transition P_buckling_, when Δε ∼Δε_c,_

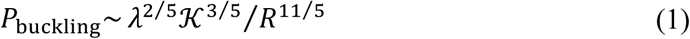

depends only on the material properties of the monolayer and on the capsule geometry, but not on the capsule stiffness *k* (Fig S6), in agreement with our experimental observations (Fig 3g, k). Above the threshold (Δε>Δε_c_), a uniform pressure P_B_ on the undeformed ring segment contribute to stabilization of the buckled ring (Fig 4d and SI Section 4).

We then tested this theory against experimental data. In order to do this, we aimed at comparing how the shape of folds depends on the pressure in the capsules with theoretical predictions. For this, we experimentally characterized the shape of single folds by their opening angle Φ, defined as the angle between the fold axis and the line connecting the lumen’s center to the epithelium detachment point, and their depth δ (Fig 4g). We reasoned that since epithelia are adhering to the capsules, experimental Φ should be comprised between two extreme values defined theoretically in absence of adhesion: α, the opening angle, and γ, the inflexion angle (Fig 4d). We find that P_B_ can be expressed in terms of α and δ, and independent of λ (Chan and McMinn, 1966) (Fig S6 and SI Section 4). Φ, δ, and P_B_ can be experimentally measured in capsules with a single fold. First, the sharp transition in experimental values of Φ and δ with time (Fig 4f) agrees with buckling being a first-order transition. Moreover, the dependence of Φ with δ is framed by the ones of α and γ with δ, fulfilling theoretical prediction with no other free parameter (Fig 4f), supporting that the folded monolayer’s shape emerges from buckling. Thus, the continuum theory of buckling correctly accounts for the shape of folds.

Next, we sought to test if our theory was accounting for the value of the buckling pressure. To test this, we estimated the elastic parameters *𝒦* and λ of the monolayer. From the relation between α^2^/δ and P_B_, we found for the bending rigidity *𝒦*∼0.5±0.2 μNμm (approximately 10^−12^ J, Fig 3i). Given that the pressure at buckling is of the order of 100 Pa (Fig 3g, l), we estimate λ≈0.1 N/m from Eq. (1) (Table 1, SI Section 5). An independent experimental measure of λ can be deduced from rigidity differences between empty versus monolayer-filled capsules measured through capsule indentation (Fig 4j and Fig S5), yielding λ=0.15±0.13 N/m (Table 1, SI Section 5) in agreement with the previous estimate. Also, previous measurements established that the Young modulus of cell monolayers was 20 kPa (Harris et al., 2012). Multiplying this value by the cell size, 10 μm, gives λ=0.2 N/m, in agreement with our findings. Altogether, the agreement between continuum theory and experimental values of elastic parameters and buckling pressure supports that folding results from buckling.

**Table 1.**
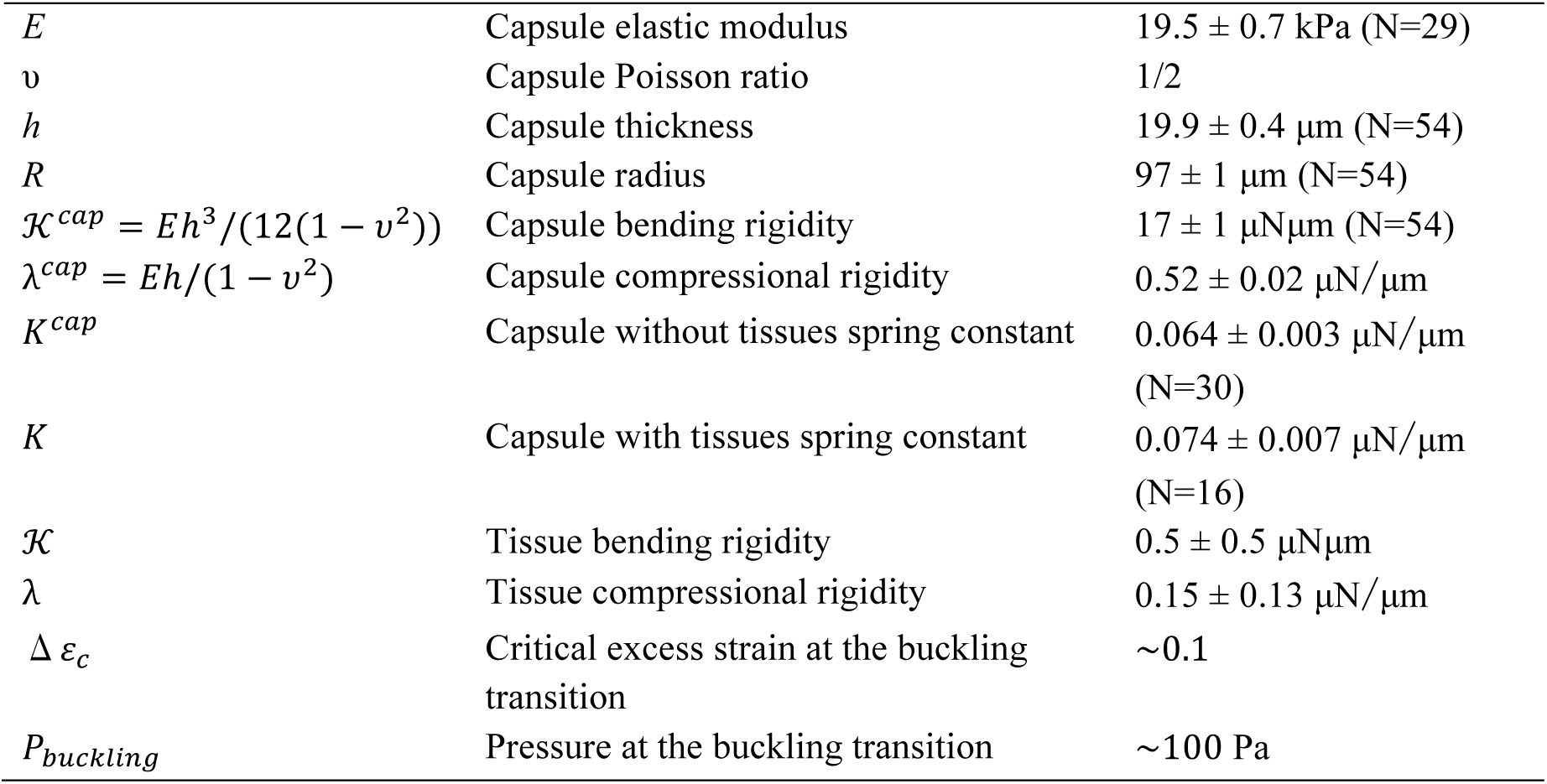
Estimation of the material parameters of the physical model. Errors are SEM.

|  |  |  |
| --- | --- | --- |
| $E$ | Capsule elastic modulus | $19.5 \pm 0.7$ kPa (N=29) |
| $\nu$ | Capsule Poisson ratio | 1/2 |
| $h$ | Capsule thickness | $19.9 \pm 0.4$ $\mu\text{m}$ (N=54) |
| $R$ | Capsule radius | $97 \pm 1$ $\mu\text{m}$ (N=54) |
| $\mathcal{K}^{cap} = Eh^3/(12(1 - \nu^2))$ | Capsule bending rigidity | $17 \pm 1$ $\mu\text{N}\mu\text{m}$ (N=54) |
| $\lambda^{cap} = Eh/(1 - \nu^2)$ | Capsule compressional rigidity | $0.52 \pm 0.02$ $\mu\text{N}/\mu\text{m}$ |
| $K^{cap}$ | Capsule without tissues spring constant | $0.064 \pm 0.003$ $\mu\text{N}/\mu\text{m}$<br>(N=30) |
| $K$ | Capsule with tissues spring constant | $0.074 \pm 0.007$ $\mu\text{N}/\mu\text{m}$<br>(N=16) |
| $\mathcal{K}$ | Tissue bending rigidity | $0.5 \pm 0.5$ $\mu\text{N}\mu\text{m}$ |
| $\lambda$ | Tissue compressional rigidity | $0.15 \pm 0.13$ $\mu\text{N}/\mu\text{m}$ |
| $\Delta \varepsilon_c$ | Critical excess strain at the buckling transition | $\sim 0.1$ |
| $P_{buckling}$ | Pressure at the buckling transition | $\sim 100$ Pa |

### The role of adhesion and proliferation in buckling studied by numerical simulations

The continuous equilibrium theory pictures well the experimental pressure and associated shapes of folds. This approach is appropriate, when dynamic processes such as proliferation and friction/detachment from the capsule shell occur on time scales that are slow in comparison with the elastic relaxation of the cell monolayer. To study the effects of these processes in a more general case, we numerically analyzed the monolayer dynamics inside alginate capsules using a 2D vertex model (Hocevar Brezavscek et al., 2012; Merzouki et al., 2016; Merzouki et al., 2017; Rauzi et al., 2015). When simulations start, cells are characterized by a resting area *A*^0^ = 300*μm*^2^ and a resting edge length *L*^0^ with (*L*^0^)^2^= *A*^0^ (Fig 5a and SI Section 6). Deviations from these values are penalized by harmonic spring energy terms with constants *K* and *k*^*s*^ for the area and the length, respectively (Fig 5a and SI Section 6) (Bruckner et al., 2017; Merzouki et al., 2016). In addition, large bending deformations of the monolayer are penalized by a harmonic spring energy term with constant *c*^*b*^ (SI). The cell elasticity *K* can be estimated by *K* ∼ λ/*A*_0_ ∼ 10^9^ N/m^3^. As in the continuum theory, the monolayer is confined to a circular domain of radius *R* by a spring constant *k* = 0.06*N*/*m* (SI Section 6). To simulate proliferation, at each iteration, one cell is selected to enter a growth phase (linear increase of *A*^0^ with time) that ends by division when the cell has increased two times its resting area. All other cells cannot enter growth or division until the growing cell has divided. We kept the other two elastic parameters *k*^*s*^ and *c*^*b*^ undefined.

**Figure 5.**
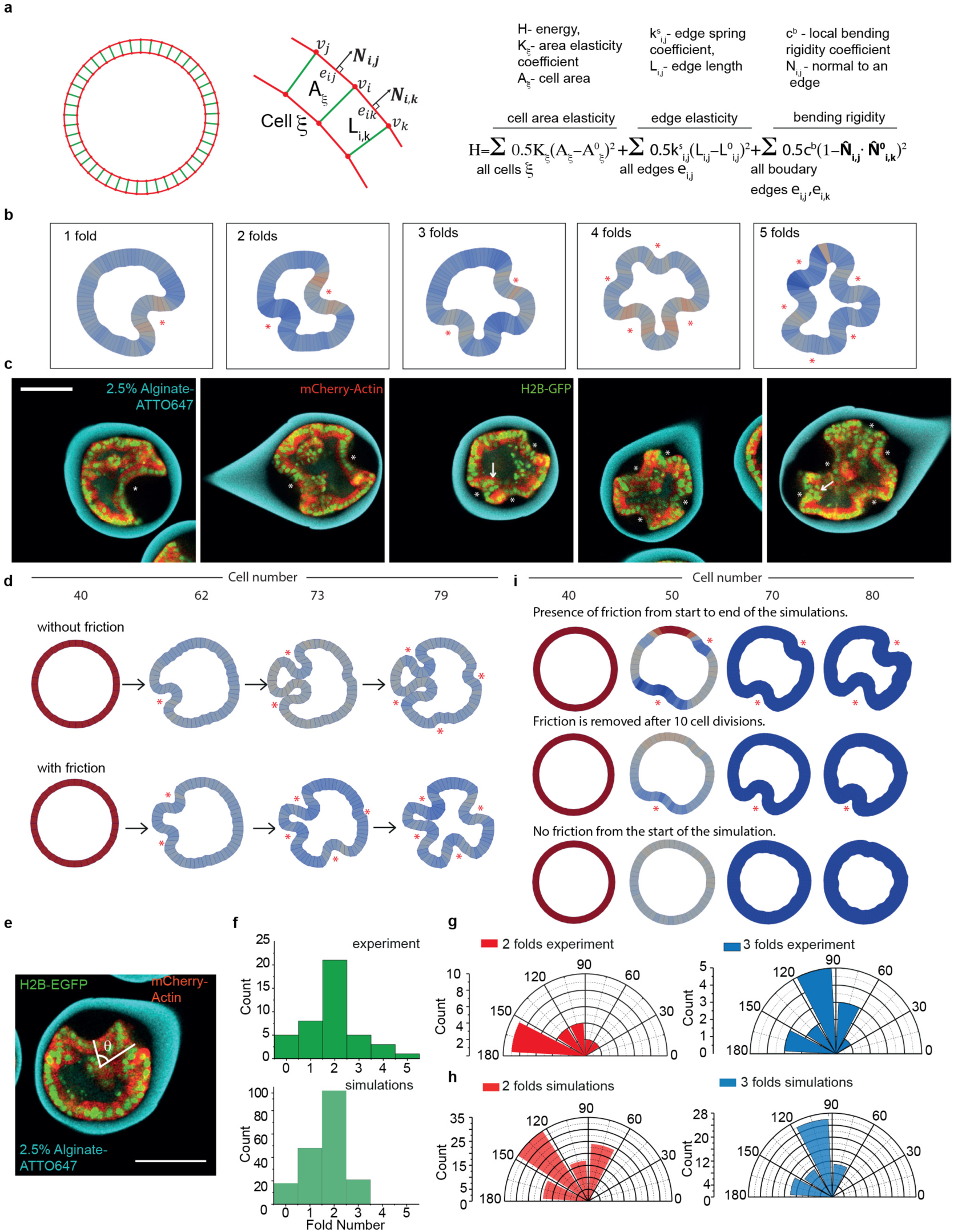
Numerical simulations of epithelial growth and buckling. **(a)** Theoretical model for numerical simulations. **(b)** Representative ending shapes of simulations executed with different couples of (*c*^*b*^,*k*^*s*^). Asterisks show folds. **(c)** Confocal equatorial planes of MDCK monolayer with 1 to 5 folds. Asterisks highlight folds. Arrows show high curved folds. Scale bar 100µm. **(d)** Top, shape evolution of a cell ring simulated without friction force. Bottom, shape evolution of the same cell ring simulated using the same sequence as on top, but with friction force *F*_*NS*_ = 5.10^−3^ *μN.* **(e)** Experimental measurement of the angle between consecutive folds, θ. **(f)** Histogram of fold number for 2.5% alginate capsules and fold number obtained *in silico* for 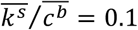. **(g)** Angle distribution between consecutive folds for 2 (red) and 3 (blue) folds in 2.5% alginate capsules. **(h)** Angle distribution between consecutive folds for 2 (red) and 3 (blue) folds obtained *in silico* for 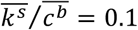 0.1. **(i)** Top, shape evolution of a cell ring simulated with friction, *F*_*NS*_ = 5.10^−3^ *μN*. Middle, shape evolution of a cell ring simulated with friction *F*_*NS*_ = 5.10^−3^ *μN.* from the start, which then got removed after 10 cell divisions. Bottom, shape evolution of a cell ring simulated without any friction.

Simulations started with 40 cells, similar to the cell number in a confocal section at confluency, and ended when the cell number doubled. Simulations reproduced folding (Fig 5b-c, Movie 6). However, at later times, the shapes of the folds differed from those observed experimentally: due to cell flows the simulated folds exhibited narrow ‘neck regions’ at their bases (Fig 5d). We reasoned that in experiments, cell adhesion to the Matrigel acts as an effective friction, that suppressed cell flows (Fig 2b) and kept the bases wide. We found in our simulations that a friction force *F*_*NS*_ ∼5.10^−3^ *μN* prevented lateral cell displacements on the capsule’s inner surface resulting in shapes resembling the experimental ones (Figs 1d & 5d, Movie 6).

Another consequence of friction was, on average, to produce more and deeper folds than without friction (Fig 5d). In experiments, two folds is the most frequent case (50%) followed by one fold (20%) (Fig 5f). This is different from the continuum theory, where equilibrium shapes feature a single fold (SI Section 3). In case of two folds, angles between folds were between 150° and 180°, whereas in case of three folds, they were between 90° and 120° (Fig 5e,g). We thought that friction could prevent mechanical relaxation away from the fold. But when friction was removed at the onset of buckling, while cells were kept proliferating, no obvious change in the shape of folds was seen even if there were less folds (Fig 5i). We suspected that proliferation could counteract the expected relaxation after removal of friction. When friction and proliferation were both removed at the onset of buckling, a clear relaxation was observed when compared to the situation where only proliferation was stopped (Fig 6a). We concluded that proliferation, by sustaining growth of the folds, and friction, by hindering propagation of the relaxation, synergistically increased the number of folds.

**Figure 6.**
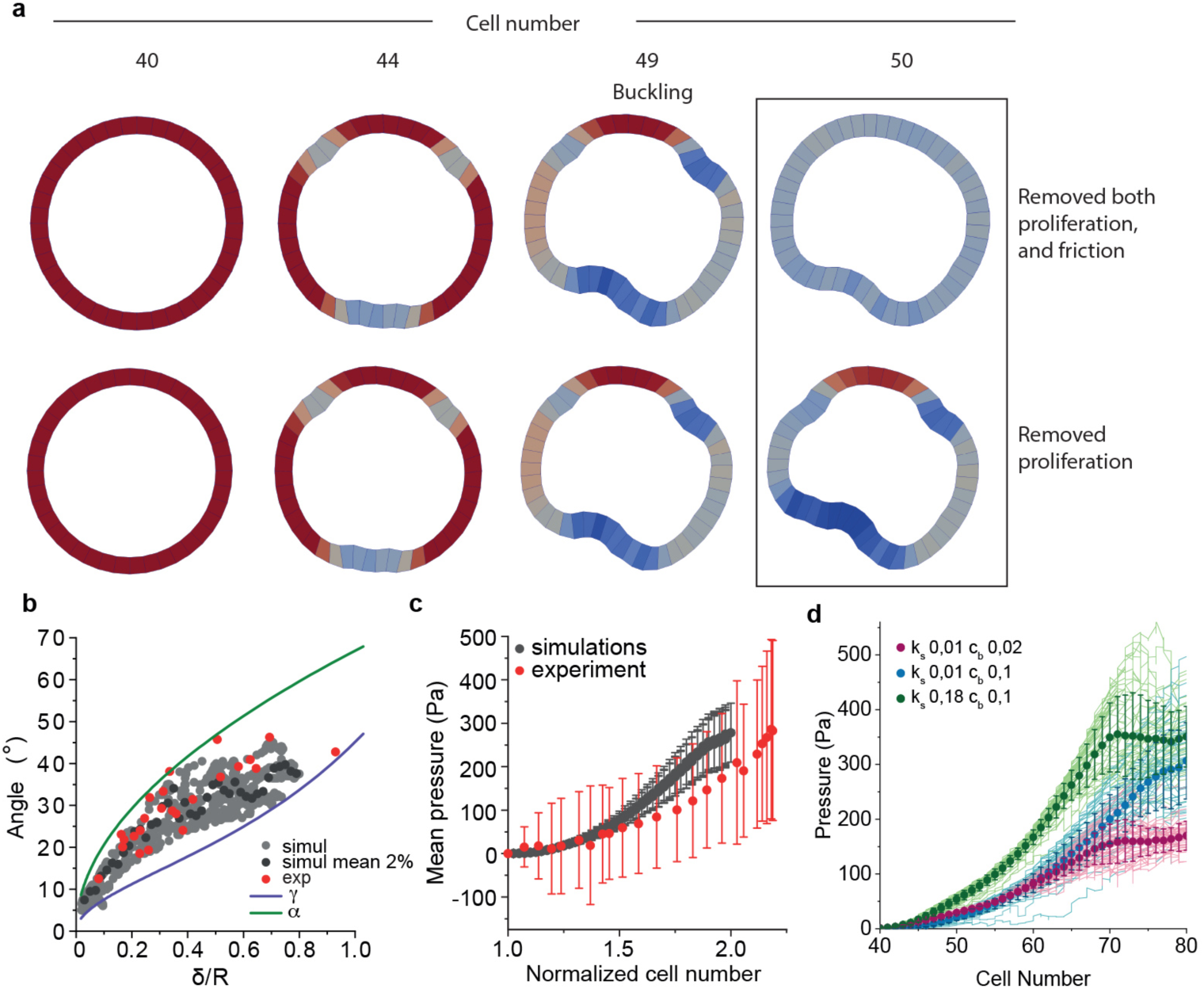
Numerical simulations of epithelial relaxation upon friction removal. **(a)** Top, shape evolution of a cell ring simulated with friction, *F*_*NS*_ = 5.10^−3^*μN*, which was removed after 10 cell divisions together with cell proliferation. Bottom, shape evolution of a cell ring simulated with friction, *F*_*NS*_ = 5.10^−3^*μN*, which was kept, however, while cell proliferation was removed after 10 cell divisions. **(b)** Angles as a function of δ/R obtained from experiments (red), continuum theory (green and purple lines) and simulations (grey and dark grey points). **(c)** Mean pressure as a function of cell number in experiments (n=53 capsules) and *in silico* (n=184); error bars are SDs. **(d)** Pressure as a function of cell number of individual numerical simulations (solid lines) and mean pressure (dots) as a function of cell number for three different pairs of normalized parameters (*c*^*b*^,*k*^*s*^).

To fix the values of the elastic parameters *k*^*s*^ and *c*^*b*^, we computed the distributions of folds for various values, and compared the distributions with the experimental distribution (Fig S7 and SI Section 7). The distribution of fold number is similar to experiments only along the line 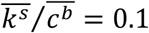, where 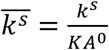 and 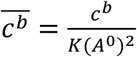 are normalized parameter used in simulations (Fig 5f and Fig S7). Furthermore, along this line, the shape of folds obtained in our simulations matched the ones observed in our experiments and continuum theory, as seen from the agreement with the relation between Φ and δ (Fig 6b).

We wondered if this set of parameters could also match the pressure dynamics in capsules. The continuum theory predicts that the pressure after buckling decreases as the excess strain increases (Fig S6). Upon a continuous increase of the excess strain (as in growing monolayers), a sharp pressure drop should be observed at the onset of folding. Instead, in experiments, the monolayer pressure typically grew continuously over time before and after buckling – only in 2% alginate capsules the pressure decreased after folding (Fig 3f). Similarly, in our simulations pressure increased monotonically after buckling for 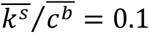 (Fig 6c-e). Away from this value, the pressure could drop after buckling (Fig S7 and SI Section 8). We thus concluded that with the specific set of parameters that matches the experimental shape, number and angular distributions of folds, we also reproduced the pressure dynamics observed in most experiments.

## Discussion

In this study, we quantitatively show that an epithelium growing under spherical confinement buckles due to the compressive stresses arising from cell proliferation. Importantly, in this case, all cells are mechanically similar, as theoretically proposed before (Hocevar Brezavscek et al., 2012; Rauzi et al., 2013). However, in previous studies, folding was obtained by differential tensions between the apical and baso-lateral sides (Hocevar Brezavscek et al., 2012), or between the lateral and the basal sides could drive the deformation (Sui et al., 2018), without the requirement of confinement and compressive forces. While our study does not exclude the importance of different contractile properties of the cells, it shows that forces required to fold the cell monolayer are in the range of micronewtons and cannot be generated by a small (<100) group of cells. As high as these forces are, the pressure necessary for epithelium buckling is at least five times lower than that required to hinder cell proliferation, which makes proliferation a potent mechanism for epithelium folding. In support of this, it was recently shown that heterogeneous growth in the drosophila wing epithelium generates buckling of proliferating cells under the confinement of surrounding tissues (Tozluoğlu et al., 2019). This supports that tissues with different growth rates can fold epithelia at different scales during organogenesis, as previously shown for the gut and the brain gyration (Savin et al., 2011; Tallinen et al., 2014).

Even if our theoretical analysis was done in 2D whereas the real system is 3D, all the main conclusion emerging for this study are valid, starting with the buckling transition being still present in 3D. Besides this, qualitative features of our description are still valid in 3D such as the presence of a single fold at equilibrium after the buckling transition or the buckling pressure being independent of the capsule rigidity. Other quantitative features should be the same, because independent of dimensionality, such as the fold’s shape and the buckling pressure. Some features are expected to change with dimensionality, such as the specific values of the vertex model parameters as well as the specific power-law relations derived in our theory.

While our study does not unravel the mechanism of epithelium folding in embryos, it identifies buckling as a potential mechanism that drives enough forces to do so. Our study also identifies the importance of friction/adhesion to the shell surrounding the tissue to promote large deformations. This is consistent with recent reports that gastrulation only completes if adhesion to the vitelline membrane is not impaired in Drosophila embryos (Bailles et al., 2019; Munster et al., 2019). Most probably, buckling will complement mesenchymal constriction (Hughes et al., 2018), apical constriction and other contractile mechanisms to drive folding in the embryo.

## Acknowledgements

Authors thank Oresti Malaspinas for his useful insights into the project.

## Author contributions

A.T. and A.R. designed the project; A.T. performed all experiments and image analyzes, with exception of cell proliferation measurements; I.D.M. helped with capsule formation, 4D confocal imaging, and performed cell proliferation measurements. K.A. and P.N. designed and fabricated microfluidic devices, and optimized the encapsulation technology. A.M. and B.C. designed numerical simulations, and A.M. performed them. C.B-M. and K.K. developed continuum theory. S.A. and J.G. measured Young’s moduli of capsules by AFM. A.T., I.D.M., C.B-M, A.M., B.C., K.K. and A.R. analyzed the results, and wrote the paper, with editions from other co-authors.

## STAR Methods

### Cell culture and generation of cell lines

Madin-Darby Canine Kidney II (MDCK-II) (ECACC, Cat. No. 00062107) cells were cultured in DMEM (Invitrogen, 10566016) supplemented with 1% (vol/vol) Penicillin-Streptomycin (Gibco BRL), 1% (vol/vol) nonessential amino acids (NEAA) 100X (Invitrogen, ref. 11140050), and 10% (vol/vol) FBS (Thermo Fisher, Cat. No.10270106) in cell culture flasks (TPP) at 37 °C and 5% CO2.

The cell line MDCK H2B-eGFP mCherry-Actin was a kind gift from the lab of Prof. Daniel J. Müller (BSSE, ETH Zurich, Switzerland). The cell line MDCK Myr-Palm-GFP, a kind gift from the lab of Dr. Mathieu Piel (Institut Curie, Paris, France), was used to generate the cell line MDCK Myr-Palm-GFP H2B-mCherry. The plasmid H2B-mCherry, a gift from Robert Benezra (Addgene plasmid # 20972), was inserted into the pLenti6.3/V5-DEST vector (containing C-terminal mCherry) using the Gateway cloning system. Lentiviral particles were generated in HEK293T cells using third generation lentiviral packaging vectors and MDCK Myr-Palm-GFP cells were infected with pLenti-H2B-mCherry. After infection, cell clones expressing both markers were sorted by Fluorescence Activated Cell Sorting (FACS) using a Beckman Coulter MoFlo Astrios, and monoclonal cells with unchanged morphology and sufficient expression level of the transgenes was selected. Cell lines were regularly tested negative for contaminationwith mycoplasma.

### Microfluidic device fabrication

The microfluidic device was printed with EnvisionTEC Micro Hi-Res Plus 3D printer using the resin HTM140V2 (EnvisionTEC), with the following printing parameters (set automatically based on the resin used): burn-in range thickness 400 μm, base plate of 300 μm, and exposure time 3000 ms. The printed device was washed using ethanol and air dried using an air gun. A thin layer of PDMS (polydimethylsiloxane) at a ratio of 1:10 (curing agent : elastomer) was put on the cone of the chip with the help of a syringe needle and baked at 70 °C for 30 minutes and subsequently baked using a UV chamber for 10 minutes. To ensure hydrophobicity and reduce the diameter of the device tip, a teflon tubing (PFA HP PLUS TUBING, 360 μm OD, 100 μm ID, IDEX) was used. The tubing was cut under a stereo binocular (Leica) using a scalpel to obtain a size of around 200 − 300 μm in length and glued on the tip of the microfluidic device with epoxyglue EA M-31CL (Loctite) and left to solidify for 1 h at RT. To make the inlets, three 19-gauge stainless steel needles were cut into segments 1: 5 cm long and polished using a Dremel 8000 WorkStation to avoid sharp edges. A small droplet of glue EA M-31CL was spread at the edges of the needles and they were inserted into the inlets of the devices, after which the glue was left to solidify for 24 hours at RT.

### Device operation and cell encapsulation

The working principle of the microfluidic device (MD) is explained in detail in (Alessandri et al., 2016). In brief, the system comprises the MD, three syringes connected to a pump (Nemesys) for flow rate control, a Matrigel cooling part, and both an Alginate charging part and a copper ring (21 mm OD, RadioSpare) connected to a High Voltage DC Power Supply (Stanford Research PS350). The MD consists of three coaxial cones inside which three different solutions are injected. The outermost cone contains alginate solution (AL), the intermediate cone contains 300 mM sorbitol solution (IS) and the innermost cone contains cells/Matrigel/sorbitol solution (CS) in a ratio of 2 : 1 : 2 (v/v), with a cell number in the range of 2*10^6^. The AL and IS solutions are loaded into two syringes controlled by the pumps for injection into the MD. The CS is injected into a cooling part to maintain Matrigel liquid, and this part is connected to a third syringe containing sorbitol that pushes out CS into the MD. The flow rates are set to 45 mL/h, 45 mL/h and 30 mL/h for AL, IS and CS, respectively, ensuring droplet formation upon exiting the MD. Once connected to the pumps the MD is positioned 50 − 60 cm above the petri dish with a 100 mM CaCl2 solution for collection of capsules. To improve capsule shape and monodispersity of size an alginate charging part and copper ring, both connected to a high voltage (2000V) generator, are introduced. The alginate charging part is a glass T connector that on opposite sides of the T has a high voltage wire (coming from the generator) and a tubing containing AL that flows down the T. The HV wire is coupled to a silver wire (OD 1 mm) that crosses the T such that it is in contact with the alginate and charges the solution, after which the charged AL then flows into the MD. The copper ring is held below the tip of the MD at a distance of 0.5 cm and centered with respect to the MD tip. The charged formed droplets passing through the copper ring under electrical tension get deflected as they cross the ring, creating a shower-like jet that prevents capsule merging. Once formed inside the calcium bath, capsules are washed with DMEM and transferred to cell culture medium.

### Fluorescent labeling of alginate

0.25 g Alginate (Protanal LF200FTS, FMS BioPolymer) was dissolved in 25 mL 0.1M MES pH 6.0 to get 1% Alginate solution. Next, 5 mg ATTO647N-amine (ATTO-TEC, ref. AD647N-95) or 13 mg Fluoresceinamine (MW 347.32 g/mol, Sigma) dissolved in 200 μL DMSO (anhydrous) was added into the tube with 25 mL 1% Alginate solution and let mix, rotating, for 10min. Next, 21.5 mg sulfo-NHS (MW 217. 14 g/mol, Fluka) dissolved in 200 μL of 0.1 M MES pH 6.0 was added and let mix for 30 min. Finally, 24 mg EDC (MW 191.7 g/mol, Sigma) dissolved in 200 μL of 0.1 M MES pH 6.0 was added and let mix and react for overnight. After this, the labeled alginates solution was transferred to Slide-A-Lyzer TM cassette 10K 12–20mL capacity and let dialyze in milliQ water for 1 day and 1 night changing the water first twice every 2 h and then last time for overnight dialysis. After dialysis the labeled alginate was filtered (Acrodisc 25 mm Syringe filter with 1 μm glass fiber media, Pall, Life Science) and keep it at 4 °C before use. The final alginate concentration was 0.5%.

To get 2.5%, 2%, 1.5% or 1% ATTO647N-labeled alginate solutions, to 10 mL milliQ water, with both 10 mL SDS 20% solution and 1 mL 0.5% ATTO647N alginate, 0.27 g, 0.165 g, 0.22 g, 0.11 g of alginate powder (Protanal LF200FTS, FMS BioPolymer) was added, respectively, and mixed overnight at room temperature. To get 2.5% FITC-labeled alginate solution, 10 mL milliQ water was mixed with 1 mL 0.5% FITC alginate, 0.25 g alginate powder and 10 μL SDS 20% solution, mixed overnight at room temperature. Before use the solutions were spun down at 48000 g for 30 min at 20 °C, after what alginate was filtered with a sterile glass fiber filter (Acrodisc 25 mm Syringe filter with 1 μm glass fiber media, Pall, Life Science).

### Imaging

To maintain capsules in set positions for several days of time-lapse acquisitions, 20 − 25 capsules were selected 24 hours post cell encapsulation and embedded in 0.4% low-melting agarose (0.04 g in 10 mL) (Sigma, 49414) in a 35 mm glass-bottom dish (Mattek, Part No. P35G-1.0-14-C). The agarose was left to solidify for 15 minutes and 2 mL of MEM containing no phenol red (Thermo Fisher, Cat. No. 51200087), supplemented with 1% (vol/vol) Penicillin-Streptomycin (Gibco BRL), 1% (vol/vol) non-essential amino acids (NEAA) 100X (Invitrogen, ref. 11140050), 10% (vol/vol) FBS (Thermo Fisher, Cat. No. 10270106) and 1% (vol/vol) GlutaMax (Gibco, Ref. 35050.038), was added. Live time-lapse confocal images of samples were obtained using inverted LSM780 NLO microscope (Carl Zeiss) using objective 20x W Plan-APOCHROMAT 20x/1.0 DIC (UV) VIS-IR (Zeiss, 421452-9800). During imaging, capsules were maintained at 37 °C with 5% CO2. For each capsule, 3D confocal Z-stacks with a range of 100 μm to 250 μm with 2 μm interval were acquired, and each capsule was imaged every 2. 5 – 3 h for 25 – 30 cycles using with definite focus (autofocus). For imaging of fixed samples, upright microscope LSM710 was used with objective 20X.

### Immunostaining

Cell monolayers in capsules were fixed with warm 4% PFA in MEM, no phenol red (not PBS, to avoid dissolving alginate capsule) for 30 min at RT. Once fixed, capsules were washed with 100 mM Glycine and 1% Gelatin in MEM. Cells were permeabilised using 1% Gelatin/0.1% Saponin in PBS 100 mM Glycine and 0.5 mM EDTA for about 45 minutes at RT until capsules dissolved followed by cell washing with 100 mM Glycine and 1% Gelatin in MEM. Cells were then incubated with primary antibodies: anti-Laminin (1 : 200, Abcam, ref: ab11575), anti-paxillin (1:250, Abcam, ref: ab26300), anti E-cadherin (1:50, BD 610181), anti-Ezrin (1:50, BD, ref: 610602), anti-p120 catenin (1:50, BD, ref: 610133) and anti-occludin (1:50, Thermo Fisher ref: 71-1500) diluted in 1X PBS 100 mM Glycine 0.5 mM EDTA overnight at 4 °C. Cell monolayers were washed 3 times with 1X PBS 100 mM Glycine and incubated with secondary antibody AlexaFluor 568 donkey anti-mouse or anti-rabbit (1:1000, Invitrogen) for 1 hour at RT (diluted in 100 mM Glycine 1X PBS with 1% Gelatin, and counterstained for f-actin and nuclei using Phalloidin488 (1:40, AlexaFluor488, Thermo Fisher, ref: A12379) and Hoechst 33342 (1:1000, Invitrogen, ref: H3570), respectively. Samples were rinsed 3 times with 1X PBS 100 mM Glycine.

### Capsule dissolution with Alginate Lyase

For high-temporal resolution experiment, cell monolayers in capsules were embedded in 0.4% agarose for imaging in a 35 mm glass-bottom dish (Mattek, Part No. P35G-1.0-14-C). The Mattek dish was priced on the side with a 19G hot needle to introduce teflon tubing into the dish sterilely for alginate lyase (Sigma-Aldrich, ref. A1603) injection. The Mattek dish was mounted onto inverted LSM780 NLO microscope (Carl Zeiss) (section Imaging). The 1mL of PBS alginate lyase solution (20 units per 1mL of PBS) was added both after 1min of imaging and after 20 min of imaging, resulting in total amount of alginate lyase of 40 units in Mattek petri dish. The imaging was performed with microscope parameters mentioned in section Imaging. For each capsule, confocal equatorial planes were acquired with time interval of 15 s for 2 hours with definite focus (autofocus). For low-temporal resolution, the experiment was conducted as a high-temporal resolution with addition of medium alginate lyase solution giving final alginate lyase amount of 75 units in Mattek dish. The imaging of capsule equatorial planes was performed with time interval of 10 min for 19 hours and with definite focus (autofocus).

### Fluorescein diffusion experiments

For fluorescein (FITC) diffusion from capsule exterior to its interior, cell monolayers in capsules were embedded in 0.4% agarose for imaging in a 35 mm glass-bottom dish (Mattek, Part No. P35G-1.0-14-C). The Mattek dish was priced on the side with a 19G hot needle to introduce teflon tubing into the dish sterilely for FITC (Sigma, ref. 46955) injection. The Mattek dish was mounted onto inverted LSM780 NLO microscope (Carl Zeiss). Once imaging started, 10 μL of 10μg/ml medium FITC solution was added. The imaging was performed with microscope parameters mentioned in section Imaging. For each capsule, confocal equatorial planes were acquired with time interval of 10 s for 2 hours with definite focus (autofocus). For FITC diffusion from capsule interior to its exterior, cell monolayers in capsules were pre incubated with FITC solution (100 μg/ml) for 2.5 hours followed by careful washing with medium. Then, cell monolayers in capsules were imbedded in 0.4% agarose for imaging. For each capsule, confocal equatorial planes were acquired with time interval of 10 min for 12 hours with definite focus (autofocus).

### Blebbistatin and Mitomycin C controls

For Blebbistatin experiments, cell monolayers in capsules were embedded in 0.4% agarose for imaging, then 10 μM of Blebbistatin (Sigma-Aldrich, ref. B0560) was added to capsules, after 1 hour of incubation at 37 °C with CO2 control, monolayers were imaged. For Mitomyocin C experiments, cell monolayers in capsules at the stage of confluence were incubated with 10 μM Mitomyocin C (Sigma-Aldrich, ref. M4287) for 1 hour followed by washing with warm medium before imaging. The imaging conditions in both cases were as described in section Imaging.

### Capsule spring coefficient measurements

There were three types of indentation experiments: i) 1% empty alginate capsule indentation, ii) 2.5% empty alginate capsule indentation and iii) 2.5% alginate capsule with a formed cellular monolayer (48 h capsule postformation). Capsules were transferred into glass-bottom Mattek petri dish with MEM containing no phenol red (Thermo Fisher, Cat. No. 51200087) solution for empty capsules and L-15 Medium (Leibovitz, Sigma, ref. L1518) supplemented with 1% (vol/vol) Penicillin-Streptomycin (Gibco BRL), 1% (vol/vol) non-essential amino acids (NEAA) 100X (Invitrogen, ref. 11140050), and 10% (vol/vol) FBS (Thermo Fisher, Cat. No.10270106) at 37 °C. The medium was changed every 2 h. Capsules were indented normally to the glass-bottom surface using micro-mechanical testing machine (FemtoTools) with indenter FT-S100. The indenter of the probe has flat silicon tip with a tip size of 50 μm by 50 μm with a force range ±100 μN and with resolution at 10 Hz 0.005 μN. For all capsule indentation experiments indentation parameters were step size 0.2 μm, speed 1 μm/s, force threshold 50 μN and signal record time 0.1 s, indentation depth was varying from 5 μm to 40 μm. The process of indentation was monitored by bright-field microscopy. To calculate capsule stiffness coefficient, the dependences of force versus indentation depth were plotted and the part of the indention was fitted with a linear fit.

### Capsule AFM indentation

Indentation measurements were performed at 37°C using a Nanowizard 4 AFM equipped with a Petri-dish heater (JPK Instruments) and an optical inverted microscope (Observer D1, Zeiss). Measurements were carried out using a PNP-TR-TL cantilevers (Nanoworld) with a nominal spring constant of 0.08 N/m after attaching a sphere of 5 μm diameter to the tip.

The force-indentation curves of Alginate capsules in concentration ranging from 1 – 2.5% were acquired then analyzed using the JPK data processing software (JPK Instruments, Germany) in which the cantilever approach curves are fitted with the Hertz and Sneddon modified model for a spherical indenter (Hertz, 1881, p. 156)

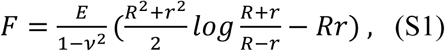

and,

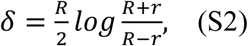

where *F* is the indentation force, δ is the indentation depth, *r* is the indenter radius, *R* is the radius of the circular contact area between indenter and sample, *v* is the Poisson’s ratio and is set to 0.5 and finally, *E* is the apparent Young’s modulus of the measured sample and is extracted from the fitted curve.

## Data Analysis

### Capsule pressure calculated from capsule deformations

To calculate the buckling pressure from capsule thinning, confocal images at capsule equatorial planes were thresholded with Fiji to get black-white masks. Then, these masks were processed using a homemade MATLAB script that detected masks boundaries (Fig. S5b), corrected for drifts (Fig. S5c), and computed outer and inner capsule radii (*R*_o_(*t, θ*)and *R*_*i*_(*t, θ*), resp.) as a function of the polar angle *θ* with the origin at the inner capsule surface centroid (Fig. S5d-e). At each polar angle *θ*, the capsule thickness

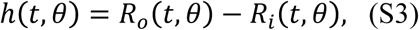

was calculated from the capsule radii, and the capsule pressure *P*(*t, θ*) was approximated as

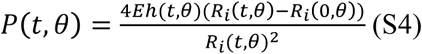

where *E* corresponds to the capsule Young’s modulus and *R*_*i*_(0, *θ*), corresponds to the capsule inner radius at the start of imaging. For spherical geometries, Eq. S4 corresponds to the theoretical relation between the inner pressure *P* and the geometrical deformations of a linear-elastic spherical shell with thickness *h*, radius *R*, Poisson ratio *v* = 0.5 and that satisfies *h*/*R* ≪ 1. For more details concerning the derivation of Eq. S4, we refer the reader to (Landau and Lifshitz, 1975, Theory of elasticity) in the main text. For instance, the reported values in Fig. S5f-g are the mean over the polar angle *θ* of the thickness given by Eq. S3 and the pressure given by Eq. S4, respectively.

### Capsule pressure calculated from capsule bending

To calculate the buckling pressure from capsule bending deformations, the capsule boundary displacement was monitored on confocal scans at capsule equatorial planes. Since alginate capsules exhibit an elastic behavior and its spring constant *k* was measured from indentation experiments, the maximal force *F*_*m*_ was calculated according to Hooke’s law: *F*_*m*_ = *kx*_*m*_, where *x*_*m*_ is a maximal capsule boundary displacement (see Fig. 3 in the main text). To convert force to pressure, we use the formula *P* = *F*_*m*_/*S*, where *S* is the surface of the deformed capsule that was approximated by a spherical cap: *S* = *2πRh*, where *R* is the radius of the spherical cap and h is the height of the spherical cap.

### Fold opening angle and fold depth calculation

Both the fold opening angle, *2α*, and fold depth *δ*, were manually measured on the confocal scans of equatorial planes of the monolayers that exhibit one-fold buckling (see Fig. 4g in the main text). The center of capsules was considered the centroid of capsule inner surface. The fold ends in contact with the capsule inner surface were considered the points where cell nuclei lied parallel to the capsule inner surface, and the angular distance between them defines the fold opening angle *2Φ* (see Fig. 4g in the main text). Fold depth *δ* was considered the maximum radial distance between the fold and the capsule inner surface (see Fig. 4g in the main text). Fold opening angle and fold depth were measured over time for each capsule. The same folds at different time points were considered as independent.

### Fold number and fold angular position calculation

Fold number was measured from the confocal scans at equatorial planes of the monolayers that exhibit developed folds. The center of capsules was considered the centroid of inner capsule surface. To analyze fold position distribution, the angles in between adjacent folds were calculated (see definition of *θ* in Fig. 5e). Angles *θ* larger than 180° were subtracted –360°.

### Code availability

The code, including a description to access and use, will be uploaded and accessible with a unique identifier code on the data repository https://zenodo.org/.

## Supplementary Figures S1-S7

**FIG.S1:**
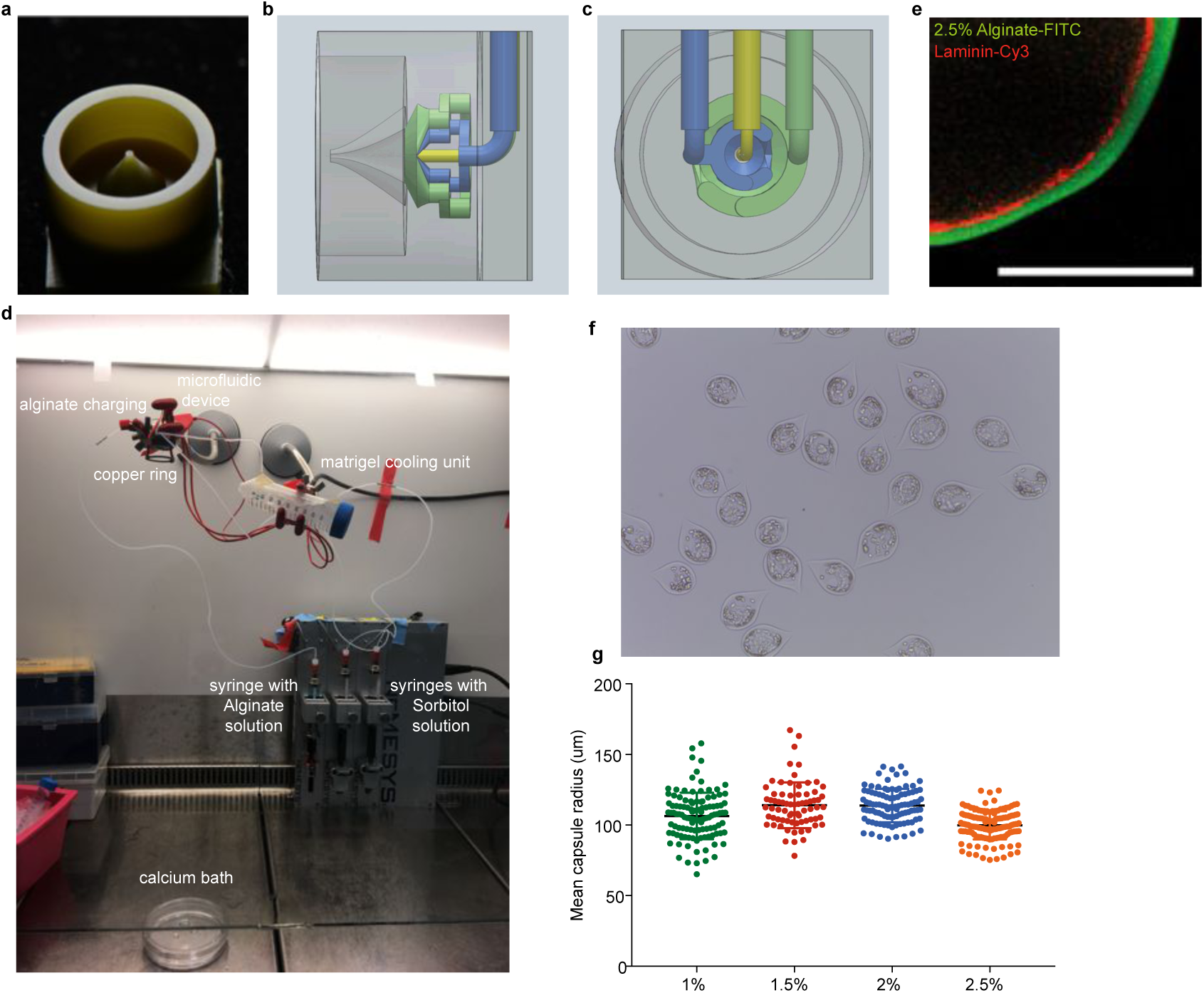
3d-printed microfluidic device and cell encapsulation experimental setup. **a**, 3D-printed microfluidic device. **b**, draft of the side view of the microfluidic device. The green, blue and yellow colors represent alginate, sorbitol solution and cell solution flows, respectively. **c**, draft of the back view of the microfluidic device. The green, blue and yellow colors represent alginate, sorbitol solution and cell solution flows, respectively. **d**, a photo of the experimental setup. **e**, confocal equatorial plane of a part of the capsule (green) showing the layer of Matrigel (red) with Cy3-labeled Laminin. **f**, Phase-contrast microscope image of 2.5% alginate capsule with growing epithelial monolayers, at 24h after capsule formation. **g**, Box-plot of mean capsule radius as a function of alginate solution percentage.

**FIG.S2:**
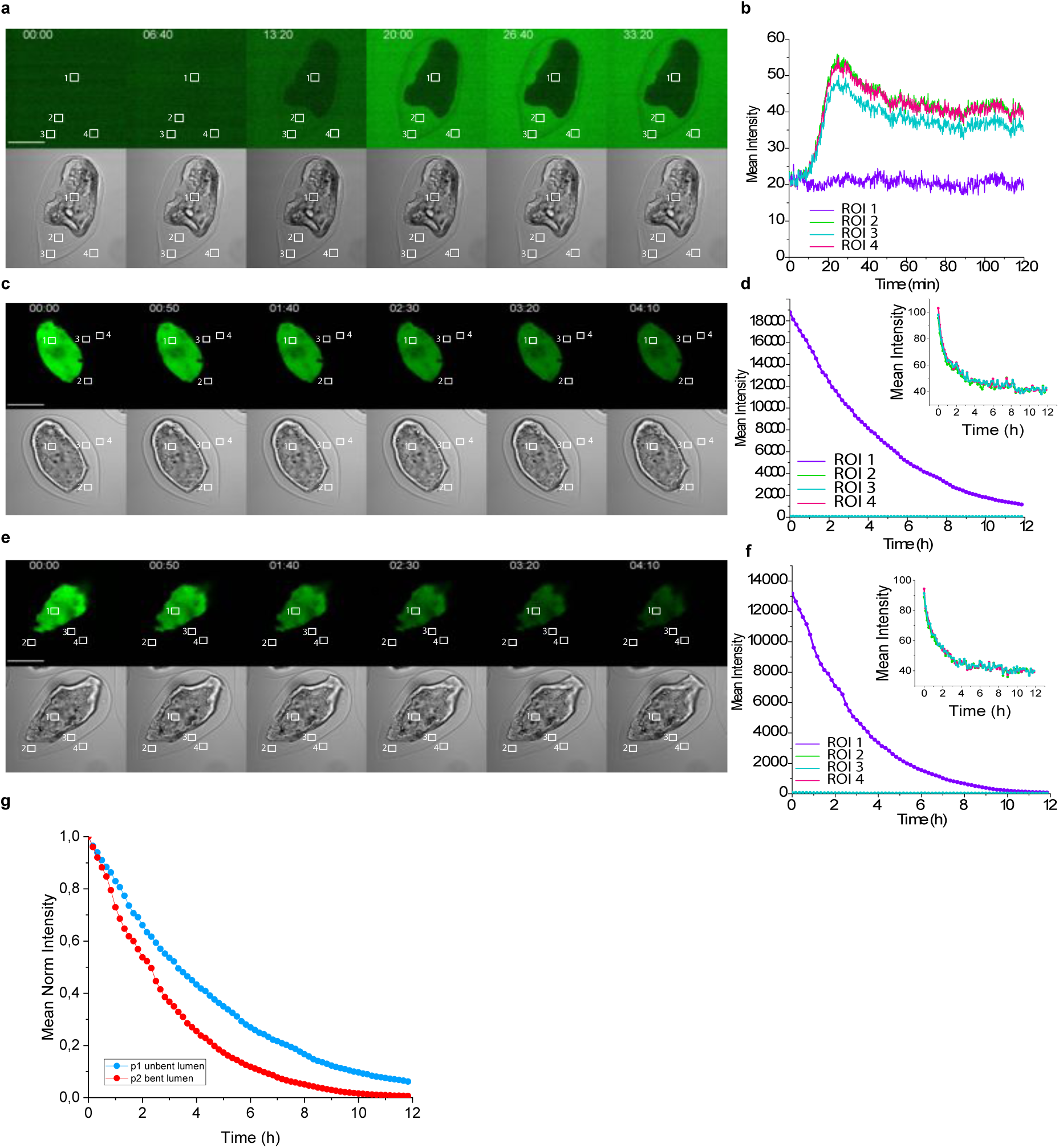
Capsule permeability and monolayer integrity controls. **a**, FITC diffusion from exterior to interior of a capsule. Top; FITC image of a confocal scan of a capsule equatorial plane containing buckled MDCK mCherry-ActinH2B-GFP monolayer. Bottom; Transmitted light image of the same capsule. **b**, Dependence of mean fluorescence intensity over time for ROIs depicted on **a. c**, FITC diffusion from interior to exterior of a capsule. Top; FITC image of a confocal scan of a capsule equatorial plane containing not yet buckled MDCK mCherry-ActinH2B-GFP monolayer. Bottom; Transmitted light image of the same capsule. **d**, Dependence of mean fluorescence intensity over time for ROIs depicted on **c. e**, FITC diffusion from interior to exterior of a capsule. Top; FITC image of a confocal scan of a capsule equatorial plane containing buckled MDCK mCherry-ActinH2B-GFP monolayer. Bottom; Transmitted light image of the same capsule. **f**, Dependence of mean fluorescence intensity over time for ROIs depicted on **e. g**, Mean normalised fluorescence intensity as a function of time for both ROIs 3 (corresponding to the ROI in the monolayer lumen) at **c** and **e**. Scale bars, 100 *μ*m.

**FIG.S3:**
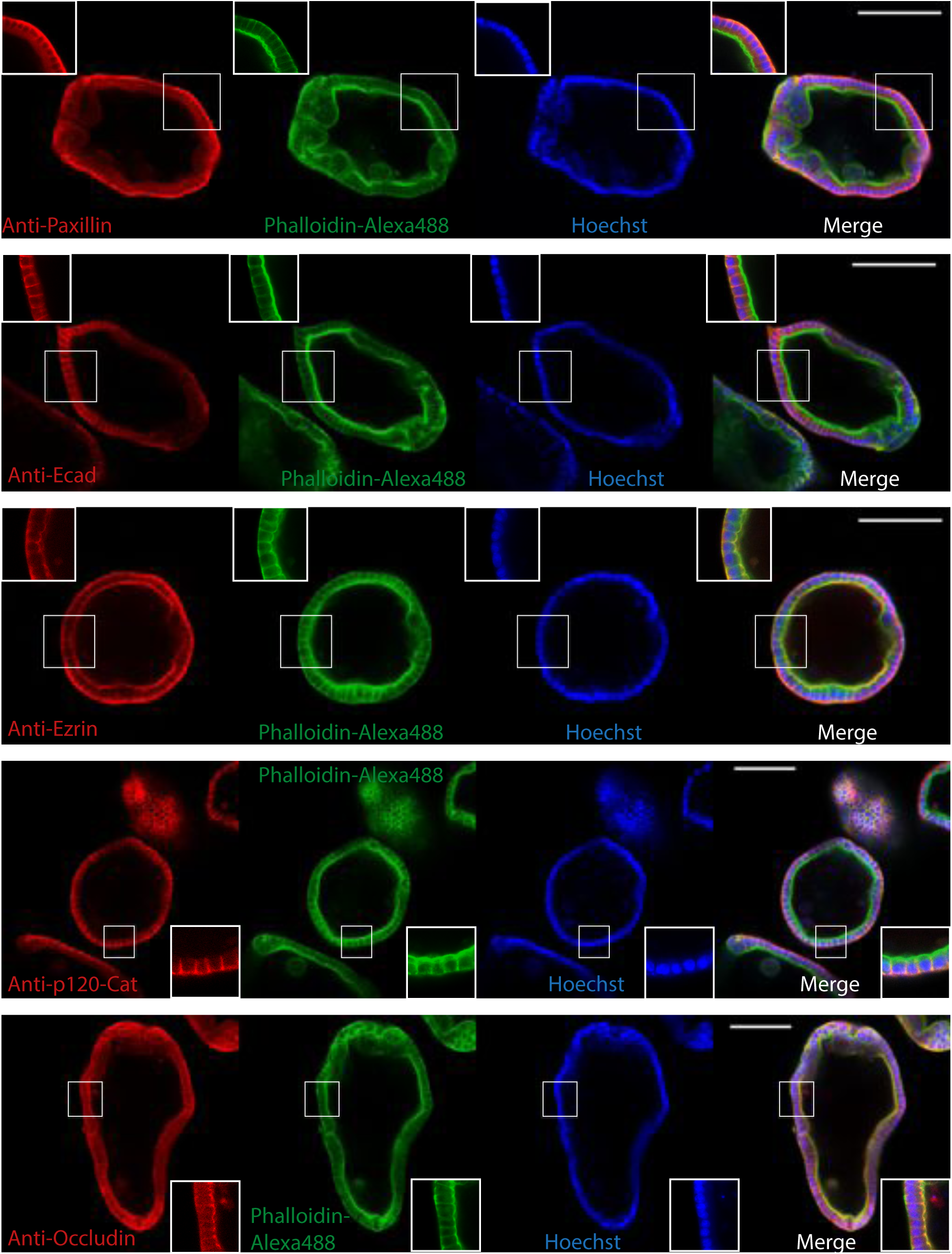
Confocal scans of immunostained formed monolayers. MDCK monolayers were fixed and immunostained to see the distribution Paxillin, Ecadherin, Ezrin, p120-Catanine and Occludin proteins. Nuclei were stained with Hoechst and filamentous actin with Phalloidin-Alexa488. Scale bars, 100 *μ*m.

**FIG.S4:**
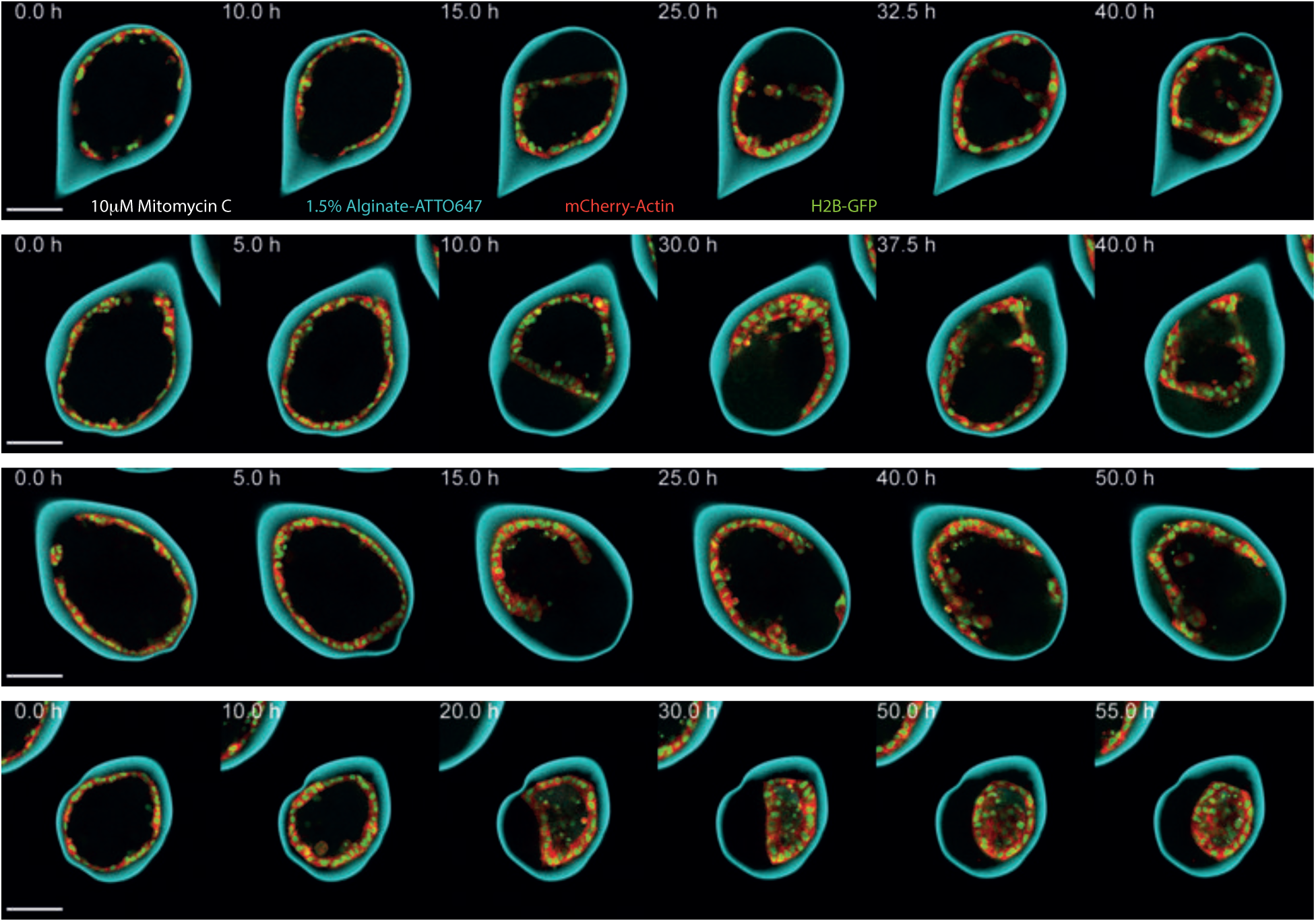
Confocal equatorial planes of MDCK mCherry-ActinH2B-GFP monolayers treated with 10 *μ*M Mitomycin-C. Scale bars, 100 *μ*m.

**FIG.S5:**
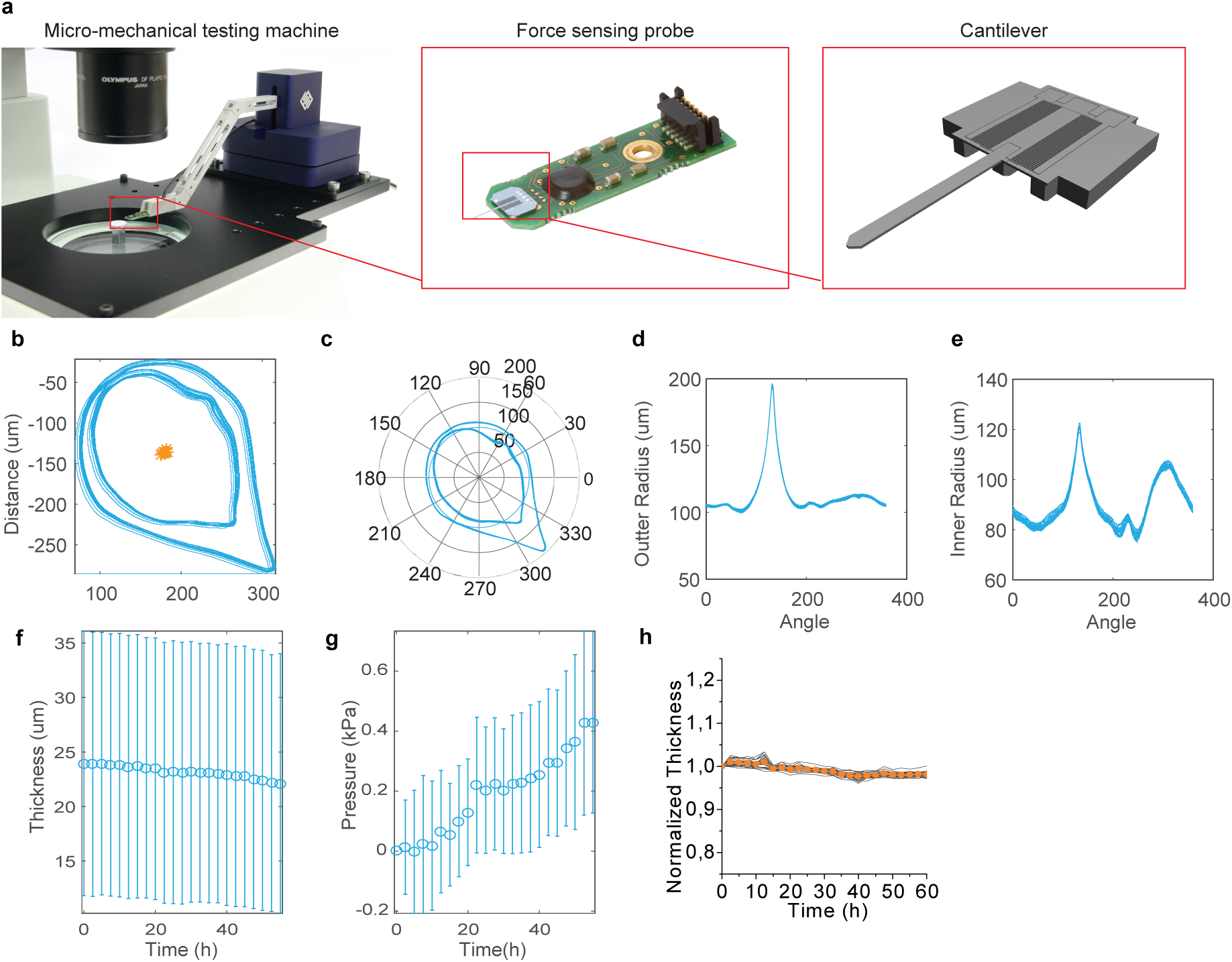
Measurement procedures of buckling pressure, fold opening angle, fold depth, and consecutive fold angle position. **a**, FemtoTools micro-mechanical testing machine. **b**, superimposed capsule contours of inner and outer boundaries corresponding to different time points. **c**, superimposed capsule contours of inner and outer boundaries corresponding to different time points in polar coordinates with the origin at capsule inner surface centroid. **d**, superimposed plots of outer capsule radii as a function of polar angle corresponding to different time points. The radii correspond to the outer contours on c. **e**, superimposed plots of inner capsule radii as a function of polar angle corresponding to different time points. The radii correspond to the inner contours on c. **f**, mean capsule thickness as a function of time for the contours in c. Each point represents a mean of 360 capsule thickness measurements corresponding to 360 angles of polar coordinates. Error bars are SDs. **g**, mean capsule pressure as a function of time for the contours in c. Each point represents a mean of 360 capsule pressures measurements corresponding to 360 angles of polar coordinates. Error bars are SDs. **h**, Normalized mean capsule thickness as a function of time for empty capsules (i.e. without cells) made of 2.5% alginate solution; n=22; error bars are SEMs. Scale bars, 100 *μ*m.

**FIG.S6:**
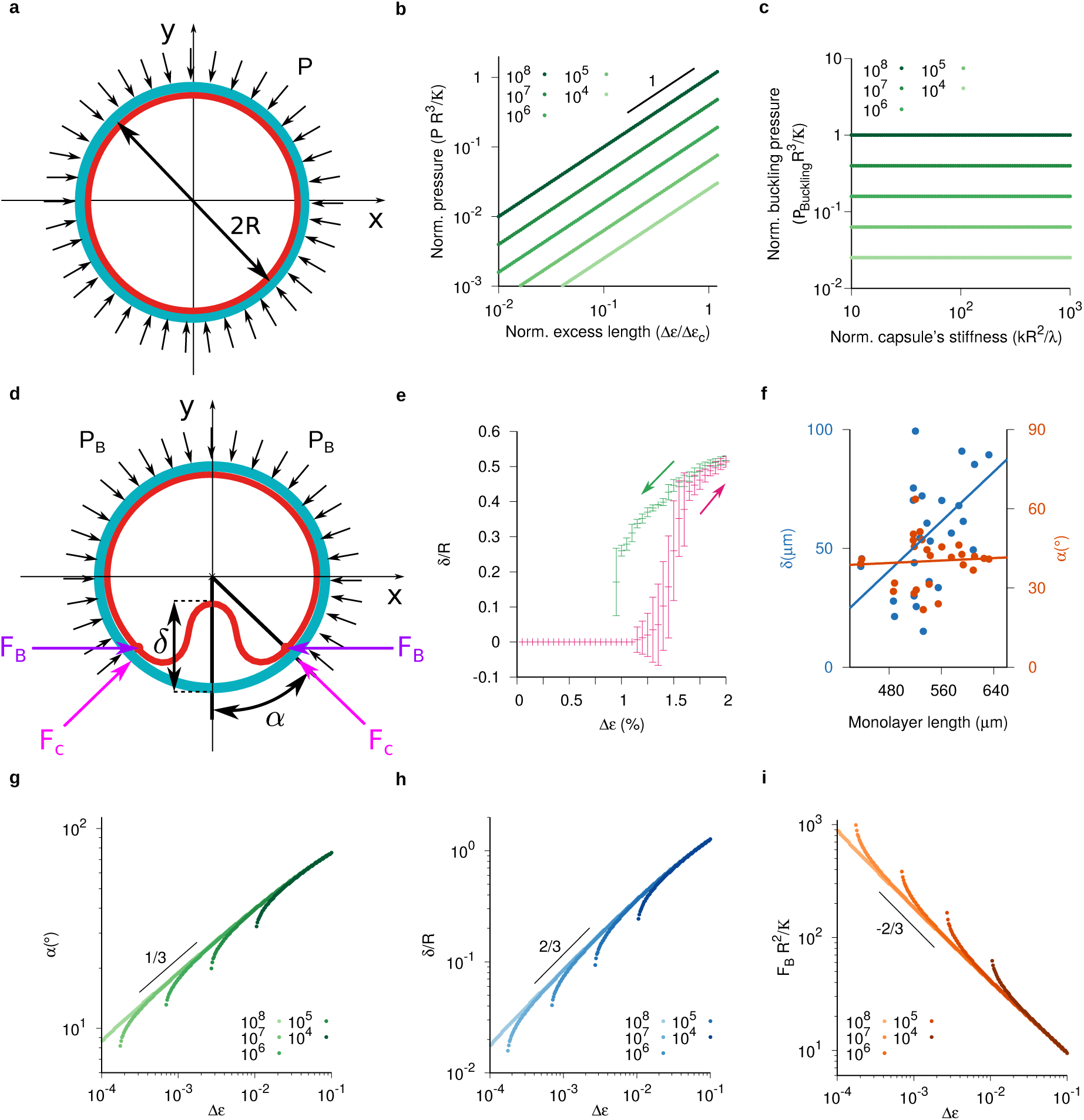
Characteristics of a confined elastic ring across the buckling instability. **a**, Schematic of a compressed elastic ring (red) under the pressure *P* of the confinement ring (cyan). *R* is the radius of the confinement. **b**, Normalized pressure (*PR*^3^*/𝒦*) as a function of the ratio between the excess strain (Δ*ϵ*) and the critical excess strain (Δ*ϵ*_*c*_) before the buckling instability (Eq. S25 for *λR*^2^*/𝒦* ≫ 1) for values of the normalized compressional rigidity (*λR*^2^*/𝒦*) between 10^4^ and 10^8^. The critical excess strain is given by Eq. S21. **c**, Normalized buckling pressure (*P*_Buckling_*R*^3^*/𝒦*) as a function of the normalized capsule stiffness (*kR*^2^*/λ*) for the same compressional rigidity values as in panel b. *P*_Buckling_ is given by Eq. S26. **d**, Schematic of a buckled elastic ring (red) under the pressure *P*_*B*_ of the confinement ring (cyan). The buckled segment is characterized by its height *δ* and its opening angle *α*. Both forces *F*_*C*_ = *P*_*B*_*R* tan (*α*) and *F*_*B*_ = *P*_*B*_*R/* cos (*α*) relate to the pressure *P*_*B*_. **e**, Fold height *δ* as a function of the excess strain Δ*ϵ*. Magenta (Green) curve corresponds to *δ* of stationary shapes for an increasing (decreasing) ramp of Δ*ϵ* from 0% (2%) to 2% (0%) at regular intervals of 0.05%. Initial conditions correspond to the stationary shape at the preceding excess strain with small amplitude fluctuations. Set of parameters: *𝒦* = 10^*-*2^, *λ* = 10^2^, *k* = 10^5^ and *R* = 1. Error bars are SD with N=30. **f**, Experimental values of *δ* and F as a function of the length of monolayers with one fold. Solid lines are linear fits. **g-i** Opening angle *α*, fold height *δ/R* and force *F*_*B*_*R*^2^*/𝒦* as a function of the excess strain Δ*ϵ*, respectively. For each panel, each curve corresponds to a distinct value of the normalized compressional rigidity (*λR*^2^*/𝒦*) between 10^4^ and 10^8^ as indicated in the legends.

**FIG.S7:**
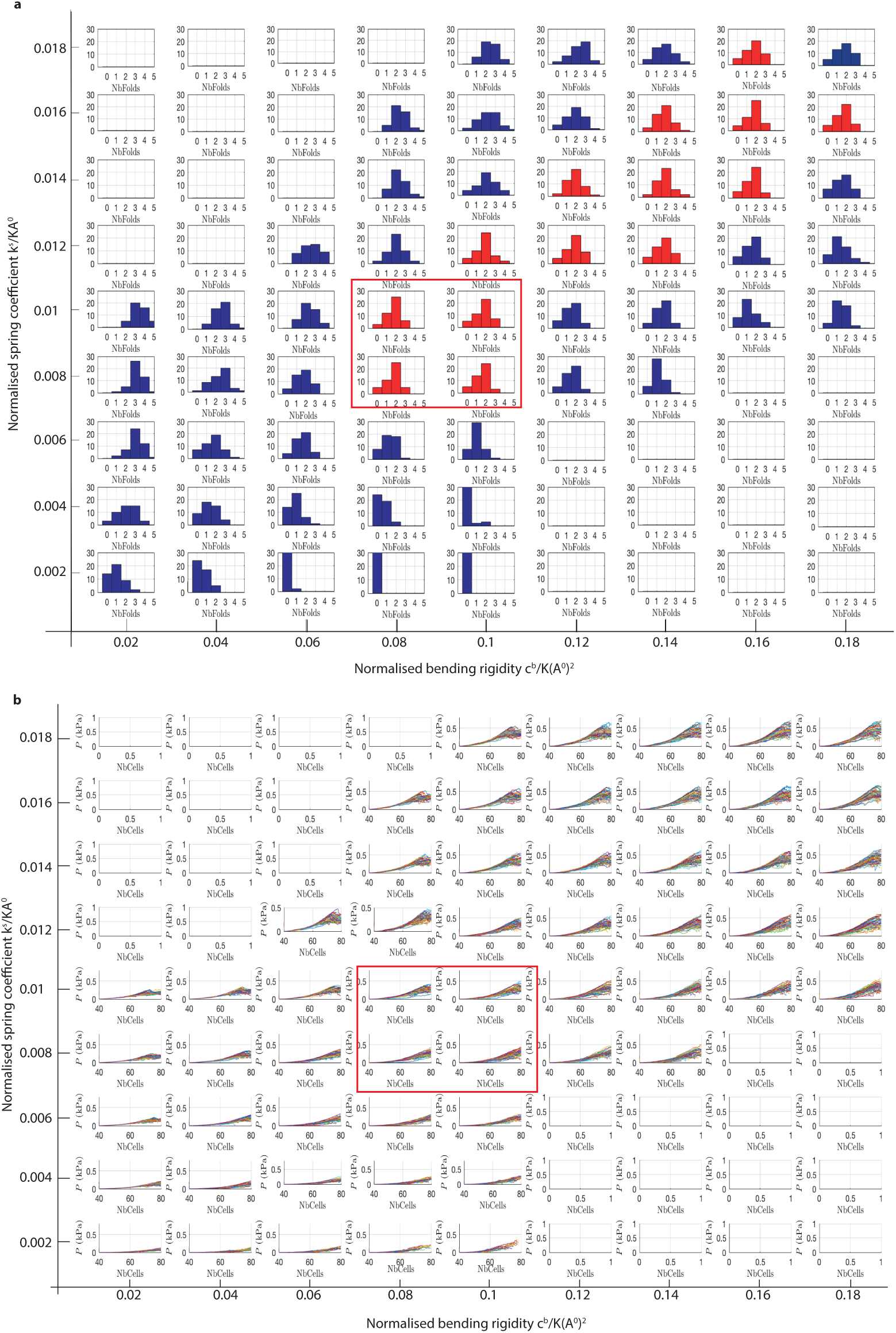
Tissue buckling within different parameter regions. **a**, Distribution of the number of folds as a function of the normalised spring coefficient *k*^*s*^*/KA*^0^ and the normalised bending rigidity *c*^*b*^*/K*(*A*^0^)^2^. The parameter regions where the distribution of number of folds match the experimental observations are coloured in red. **b**, Pressure evolution within different parameter regions. Evolution over time (the number of cells increase from 40 to 80) of the pressure *P* applied by the growing tissue on the elastic capsule within different parameter regions (*k*^*s*^*/KA*^0^ and *c*^*b*^*/K*(*A*^0^)^2^). In both panels, the red square marks the parameters regime that fits best experiments.

## Supplementary text

### Section 1: Estimation of the water flux across a monolayer upon folding

In the following, we estimate the flux of small molecules across a monolayer and compare its value with measurements for epithelial monolayers.

The volume of a monolayer lumen (i.e. hollow space enclosed by the monolayer) is estimated by considering that a monolayer lumen is a sphere with a radius *R* ∼ 100 *μ*m. Hence its volume is of the order of *V* ∼ 4·10^−12^ m^3^. Similarly, the area of a monolayer lumen can be estimated by assuming a spherical geometry with the same characteristics as above, yielding an area of the order of *A* ∼ 10^−7^ m^2^. Next, we consider that over the *T* ∼ 2 h ∼ 7000 s that folding lasts, a monolayer lumen loses half of its volume due to a water outflow *J* across the surface of the monolayer lumen. Therefore, this water outflow is of the order of

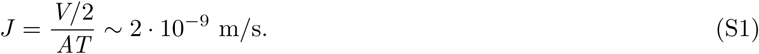

Of note, the premise that the total volume lost is half of the initial volume is in general an overestimation, according to our experiments. Previous measurements of small-molecule fluxes across epithelial monolayers in other contexts range between 10^−6^ – 10^−8^ m/s, depending on the molecule type (Rosenthal et al, 2019, p. e13334; Rosenthal et al, 2010, pp. 1913-1921). This suggests that water evacuation from the lumen to the exterior is not hindering the kinetics of monolayer folding.

### Section 2: Description of a confined elastic ring (macroscopic approach)

In this section, we introduce a description for the mechanics of epithelial monolayers weakly adhered to the inner surface of an elastic spherical shell. In particular, we focus our attention on the understanding of the tissue dynamics near the transition between a spherical state into a buckled state.

Alginate capsules act as rigid containers that constrain the growth of tissues. To a good approximation in our setting (see main text), alginate capsules are regarded as thin linear elastic materials with a Young’s modulus *E* ∼ 20 kPa, a thickness *h* ∼ 20 *μ*m and a Poisson ratio *ν* = 1/2. Hence, in the thin-layer limit, alginate capsules are akin to elastic shells with bending rigidity *𝒦*^*cap*^ = *Eh*^3^/12(1 – *ν*^2^) and compressional rigidity *λ*^*cap*^ = *Eh/*(1 – *ν*^2^) (Landau and Lifshitz, 1975, Theory of elasticity). These material parameters are of the order of *𝒦*^*cap*^ = 20 *μ*N·*μ*m and *λ*^*cap*^ = 0.5 *μ*N/*μ*m, according to the parameter set in Table S1.

Epithelial monolayers are active materials with the capacity to spontaneously undergo morphological changes regulated by cell forces. When confined to flat surfaces, epithelial cells often reach a confluent state in which the cell density plateaus and cell motion is drastically reduced, similar to glass phases of dense active particle suspensions. Since the experimental analysis is done for a confocal slice, we describe the cell layer as a cylindrical shell and assume translational invariance along the cylinder axis. Unless otherwise stated, we consider only the cross section of the cylinder and look at deformations of a ring. The lack of cell flows at the onset of the buckling transition suggests that the complex mechanics of encapsulated epithelial monolayers can be approximated by a 2*d* thin elastic ring. Unlike alginate capsules, their bending rigidity *𝒦* and compressional rigidity *λ* are regarded as independent material parameters a priori, which might be regulated through cell-cell adhesion, cell-matrigel adhesion or cell contractility. Therefore the global energy per unit length of the 2*d* system constituted by the monolayer and the capsule reads

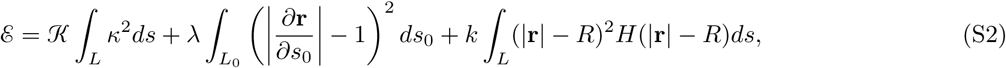

The first term represents bending energy with *κ* being the local mean curvature in 2*d*. The second term represents the strain energy where the integrand stands for the relative displacement between two material points with respect to their equilibrium configuration. The third term represents the adhesion-free interaction between the tissue and the capsule. For small capsule deformations, this interaction is approximated by a piece-wise harmonic potential with a spring constant *k* dependent on the material properties of the capsule, where *H*(|**r**| **–** *R*) is a Heaviside step function and *R* is the radius of the alginate capsules (see Fig. 4 in the main text). The current elastic-ring shape is parametrized by the arc-length *s* with its length being *L*, whilst the equilibrium elastic-ring shape in the absence of constraints is parametrized by the arc-length *s*_0_ with its length being *L*_0_.

The area of the monolayer is found experimentally to increase over time as a result of cell proliferation. The observed linear growth in cell number yields a doubling time scale of about ∼ 60 hours. Remarkably, this time scale is significantly larger than the time scale of buckling formation ∼ 10 hours, suggesting that *L*_0_ variations can be ignored during tissue folding dynamics as a first order approximation. The separation of these two time scales enables to decouple the dynamics driven by cell division to the mechanical relaxation of cellular forces. Thereby, as a simplifying approximation we assume that the system equilibrates before large density variations occur and presume that the resting length *L*_0_ is a time-independent parameter. The effects induced by the competition between cell proliferation and cell mechanics relaxation time scales is discussed elsewhere. Therefore taking as unit length *R* and as energy unit *𝒦/R*, the mechanics of elastic-ring folding are fully characterized by three dimensionless parameters: the elastic-ring compressional rigidity *λR*^2^*/𝒦*, the capsule spring constant *kR*^4^*/𝒦* and the ratio between the elastic-ring length and the capsule length *L*_0_/2*πR*.

#### Numerical methods

The details of the numerical method employed to study the statistical properties of our system are as follows: at a given instant of time, the state of an elastic ring is described by the position of *N* material points which are located at positions **r**_*i*_ = (*x*_*i*_, *y*_*i*_) with *i* ∈ (0,*N –* 1). Typically, *N* = 100. The energy per unit length assigned to a certain configuration is given by the discretized form of the mechanical energy (S2), namely

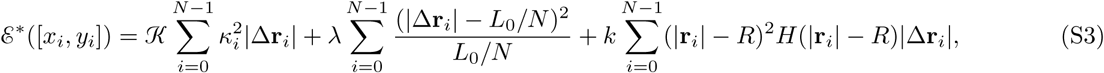

where 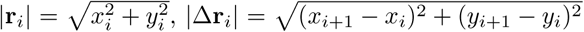 and *κ*_*i*_ = 4((*y*_*i*+1_ − 2*y*_*i*_ + *y*_*i*−1_)(*x*_*i*+1_ − *x*_*i*−1_) − (*x*_*i*+1_ − 2*x*_*i*_ + *x*_*i*−1_)(*y*_*i*+1_ − *y*_*i*−1_))/((*x*_*i*+1_ − *x*_*i*−1_)^2^ + (*y*_*i*+1_ − *y*_*i*−1_)^2^)^3/2^ is the curvature at the i-th position. Because of the ring geometry *x*_*N*_ = *x*_0_ and *y*_*N*_ = *y*_0_.

To compute the configurations with minimal energy, we introduce an artificial dissipative dynamics that drives the system toward a local minimum of the energy (S3). For each material point dynamics is determined by

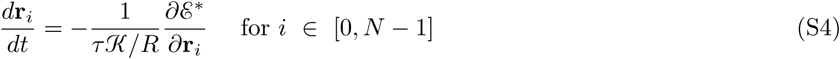

For us, the time scale *τ* is arbitrary and thus, it is set to *τ* = 1. Notice that the system might be trapped in metastable states in the absence of bulk fluctuations.

### Section 3: Buckling instability induced by confinement elasticity

In the present section, we revisit the problem of an inextensible elastic ring introduced inside an undeformable rigid circular ring. This problem has been investigated to various degrees and in different ways by a number of researchers: see for instance (Lo et al, 1962, pp. 691-695; Chan and McMinn, 1966, pp. 433-442) in the main text.

For simplicity, we focus first in the limit of inextensible rings and underformable confinements, meaning that *λ/𝒦R*^2^, *k/𝒦R*^4^ ≫ 1 in Eq. S2. Thereby the elastic-ring forces are neither sufficient to deform capsules nor to compress/extend material elements (i.e. *L* = *L*_0_). We solve numerically Eq. S3 to obtain the equilibrium shapes. When the monolayer length is smaller than the length of the capsule (*L*_0_/2*πR* ≤ 1), the elastic ring attains a circular shape regardless of the confinement as there are no contact interactions between them. When the ring length is larger than the length of the confinement (*L*_0_/2*πR >* 1), however a buckling instability occurs. Note that elastics rings attain a shape with a single fold induced by geometrical confinement.

To investigate further the emergence of a buckled ring shape above a certain length *L*_0_, we study analytically the behaviour of elastic rings near the transition point. To this end, we propose to account for the morphological variations as the system undergoes the buckling transition by the following parametrization for the ring shape

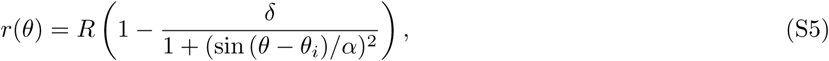

where *r* and *θ* are the radial and angular coordinate with respect to the confinement center of mass, respectively. In cartesian coordinates, the shape curve is parametrized as (*x*(*θ*), *y*(*θ*)) = (*r*(*θ*) cos (*θ*), *r*(*θ*) sin (*θ*)). For *δ ≠* 0, the ring exhibits a folded region located at the random angular position *θ*_*i*_. Its shape is characterized by two coefficients: *δ* that controls the fold amplitude and *α* that controls its angular width.

The length of an inextensible ring parametrized by Eq. S5

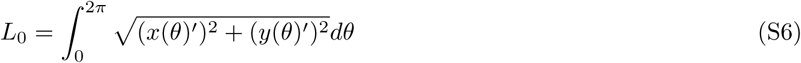

is a conserved quantity. Considering that the coefficients *δ, α* ≪ 1, the leading order corrections in *δ* and *α* of Eq. S6 can be recast as

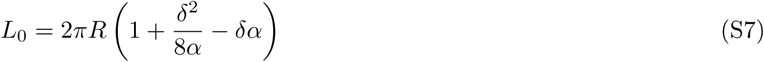

By defining the excess strain Δ*ϵ* = Δ*L/*2*πR* as the excess length Δ*L* = *L*_0_ − 2*πR* of the ring with respect to the confinement perimeter normalized by the latter, Eq. S7 can be expressed as

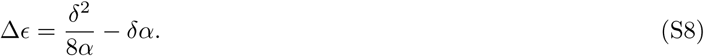

Likewise, the bending energy of an inextensible ring parametrised by Eq. S5 is expressed as

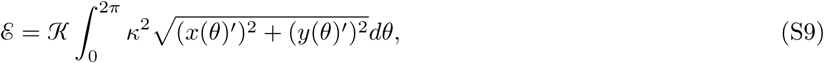

where the local curvature reads *κ* = (*y*(*θ*)″ *x*(*θ*)′ − *y*(*θ*)′*x*(*θ*)″)/((*x*(*θ*)′)^2^ + (*y*(*θ*)′)^2^)^3/2^. Up to leading order corrections in *δ* and *α*, Eq. S9 can be recast as

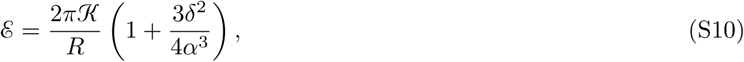

Combining Eqs. S8, S10 and calculating the shape that minimizes the energy (S10) with the geometrical constraint given by Eq. S8, we deduce power-law relations between the geometrical properties of the buckled ring (*δ, α*) and its current excess strain Δ*ϵ*, which read

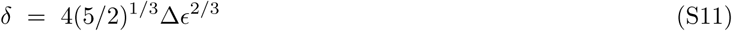

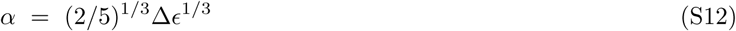

As Δ*ϵ* increases progressively beyond the value at the buckling transition, both parametric coefficients increase but the relative variation of them is different. We predict that for the same variation in *L*_0_, the amplitude of the fold *δ* increases more than the angular width *α*, thereby conditioning tissue growth.

The sub-linear variations of the energy per unit length of the minimal buckled shape against the excess strain read

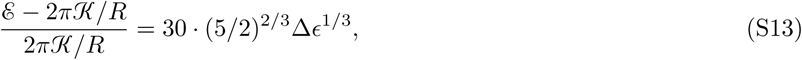

#### Equilibrium number of folds at the buckling instability

Here we present an argument to show that the minimal shape near the buckling transition exhibits at most one prominent fold. To address this question, we follow a similar argument as above and parametrize a ring shape with *n* equal folds by the function

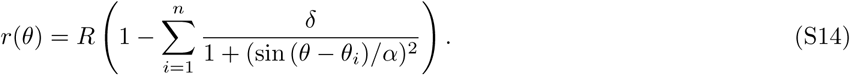

We ignore geometrical heterogeneities among folds as presumably they would lead to configurations with overall higher mechanical energies. Remember that *r* and *θ* are the radial and angular coordinate with respect to the capsule’s geometrical center, respectively. For *δ ≠* 0, the shape is buckled *n* times at the angular positions *θ*_1_, *…, θ*_*n*_ with the distance between two consecutive folds *θ*_*i*+1_ − *θ*_*i*_ = 2*πR/n* being constant.

As we introduced above, when the excess strain Δ*ϵ* = (*L*_0_ − 2*πR*)/2*πR* ≪ 1, the deformations of the ring shape are expected to be small as well, meaning that both parametric coefficients *δ, α* ≪ 1. In this situation, we can cast the leading corrections of the length *L*_0_, which is constant for inextensible elastic rings, as follows

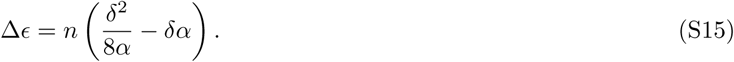

Notice that the excess strain of a ring with *n* folds is simply given by the excess strain of a single fold (S8) times the number of folds *n*, as expected from mean-field considerations. Similarly, the correction to the total energy per unit length (S2) near the buckling transition also scale linearly with the number of folds and it reads

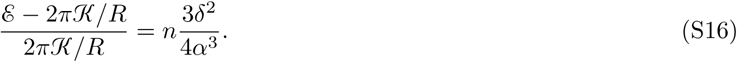

Combining Eqs. S15-S16 and calculating the minimum of (S16) with the geometrical constraint given by Eq. S15, we deduce the power-law relations between the geometrical properties of the buckled ring (*δ, α*) and its excess strain Δ*ϵ* and number of folds *n*, which read

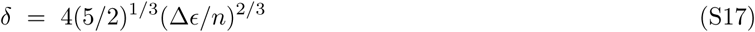

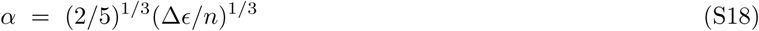

In contrast to the ring length *L*_0_ Eq. S15, the geometrical properties of the fold are not proportional to the number of folds. Both parametric coefficients follows a power law with the number of folds *n* and the exponent is smaller than 1. From the expression of *δ* and *α*, the energy per unit length of the minimal buckled shapes becomes

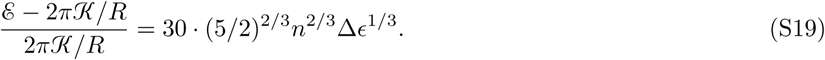

Remarkably, the energy increases with the number of folds *n* in a sublinear manner. As a result, the shape with overall minimal energy has one fold for small excess strain Δ*ϵ* ≪ 1.

#### Asymptotic limits of the buckling transition for confined elastic rings

In the sections above, we focus our study in the ideal limit of an inextensible elastic ring and rigid capsules to understand the mechanics of buckled tissues under confinement. However, the compressibility of epithelial cells can not be disregarded in our setting, as prior to the buckling transition, cell shape variations are observed. These observations suggest that cells have a finite compressional rigidity *λ* whose value is estimated from experimental measurements of capsule elastic deformations (see Table S1). In the present section, we explore how the buckling transition is influenced by the capsule and tissue rigidities.

In the general situation, two limiting cases can be distinguished for elastic-ring deformations controlled by the rigidity of capsules. When comparing the energetic contribution attributed to elastic-ring and capsule deformations in Eq. S2, a dimensionless parameter emerges *λ/kR*^2^. Hence, the tissue growth is influenced by the ratio between the cellular compressional rigidity *λ* and the stiffness of the capsule *k*. Notice that the spring constant *k* relates to the material properties of the capsule such that *k* = *λ*^*cap*^*/R*^2^, as it is discussed in the section below. In consequence, the dimensionless parameter *λ/kR*^2^ is independent of the capsule radius *R*, but not of capsule and cell mechanics.

For *λ/kR*^2^ ≪ 1, the deformations of cell shapes are energetically favorable against capsule deformations, meaning that below the buckling transition, cells tend to accommodate the internal stress generated by cell proliferation through shape deformations. As a consequence, when cells are compliant, the threshold of the buckling transition is set by balancing bending and compressional cell deformations. In particular, for *λR*^2^*/𝒦* ≪ 1 cells are infinitely compressible, so that the excess strain Δ*ϵ* is fully accommodated through cell compression. In this situation, the minimal shape of the elastic ring is a circle of radius *R* and the leading correction of the total energy per unit length as a function of the small excess strain Δ*ϵ* = (*L*_0_ − 2*πR*)/2*πR* ≪ 1 reads

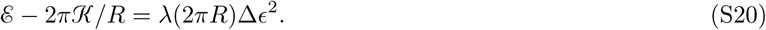

Alternatively, the limit of inextensible elastic rings (i.e. *λR*^2^*/𝒦* ≫ 1) yields an energy per unit length of the minimal buckled ring that is given by Eq. S13. The boundary between these two regimes can be delimited by comparing both energies (S13) and (S20), setting a critical excess strain

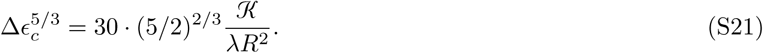

Next, we further investigate the influence of tissue elastic parameters on the buckling transition. We numerically compute minimal shapes as a function of the excess strain Δ*ϵ* and the dimensionless parameter *𝒦/λR*^2^. Fig. 4e in the main text shows two distinct morphological patterns for elastic rings, corresponding to circular rings and buckled rings. The critical excess strain Δ*ϵ*_*c*_ given by Eq. (S21) is in qualitative agreement with numerical simulations. Consequently, when Δ*ϵ <* Δ*ϵ*_*c*_, the ring attains a compressed circular shape of radius *R*, but for Δ*ϵ >* Δ*ϵ*_*c*_, the ring buckles.

Next, we investigate the nature of the buckling transition by studying the stability of minimal shapes of Eq. S2 near the transition point. To this end, we numerically compute minimal shapes as a function of the excess strain Δ*ϵ* for a fixed set of elastic parameters. The excess strain Δ*ϵ*, first increases from 0 to 0.02 and second decreases from 0.02 to 0 at regular intervals of 0.0005. At a certain Δ*ϵ*, the initial condition corresponds to the stationary shape at the preceding excess strain plus small amplitude fluctuations. Fig. S5e shows the height of the folded region of an elastic ring *δ* as a function of the excess strain, being *δ* = 0 for circular rings and *δ ≠* 0 for buckled rings. We observe signatures of hysteresis that is characteristic of first order transitions. The threshold for the increasing ramp of Δ*ϵ* is not coincident with the threshold on the decreasing ramp, delimiting a coexistence region for intermediate excess strains, that in addition is manifested by larger error bars. We checked that the hysteresis region is robust to variations in the elastic parameters. Therefore, we conclude that the nature of the buckling transition is first order.

For *λ/kR*^2^ ≫ 1, the deformations of the capsule are energetically favourable against cell deformations, meaning that below the buckling transition, cells tend to expand and deform the capsule to accommodate the forces generated by cell proliferation. As a consequence, when capsules are compliant, the threshold of the buckling transition is set by balancing cell bending deformations and capsule compressional deformations. In particular, for *kR*^4^*/𝒦* ≪ 1 the capsules are infinitely compressible, so that the excess strain Δ*ϵ* is fully accommodated through capsule inflation. In this situation, the minimal shape of the elastic ring is a circle of radius *L*_0_/2*π > R* and the leading correction of the total energy per unit length as a function of the small excess strain Δ*ϵ* = (*L*_0_ − 2*πR*)/2*πR* ≪ 1 reads

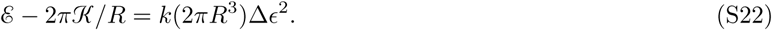

In a similar spirit as above, the energy per unit length given by Eq. S13 for an inextensible ring inside an undeformable capsule can be employed to estimate the boundary between these two limiting cases, which sets the critical value for the excess length

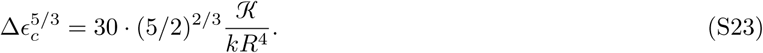

Consequently, when Δ*ϵ <* Δ*ϵ*_*c*_, the ring attains a circular shape of radius *L*_0_/2*π > R*, but for Δ*ϵ >* Δ*ϵ*_*c*_, the ring buckles.

### Section 4: Forces of the ring-confinement system

In our setting, we can investigate the interplay between spherical alginate capsules and the morphological changes of epithelial tissues, and how these in turn are driven by the mechanical forces generated at the interface between capsules and cells. Indeed, epithelial monolayers can build up large-scale collective forces mediated by cell-cell contacts. These forces suffice to induce large deformations on capsules with elastic modulus of ∼ 10 kPa. In the course of tissue buckling, distinct modes of deformations are identified. In the present section, we summarize the analytical arguments used to estimate the tissue forces from capsule deformations.

#### Isotropic pressure of an elastic ring before buckling

At the early stages, cells migrate freely on the inner surface of alginate capsules. Upon reaching a confluent state, in which no free spaces remain across the monolayer, cells keep on replicating and thus generating local force dipole at each of these events. The equilibration of these forces at the tissue-scale lead to an overall cellular compression, evidenced by the observed progressive increase of cell height or decrease of cell in-plane area. These compressive forces are transmitted to the substrate underneath, resulting in a measurable expansion of the alginate capsule. This first phase terminates, when the tissue folds.

To investigate the nature of the forces before buckling, let us consider a 3*d* elastic shell of radius *R* + *δR > R* with *δR* ≪ 1 that is confined in a shell of smaller radius *R* under the action of a uniform pressure *P*, Fig. S6a. The departure of the capsule from its equilibrium state is balanced by the work done by an external pressure *P* Fig. S6a. The variation of the curvature in the compressed state is ∼ *δR/R*^2^, and the total bending energy scales as *𝒦* (*δR/R*^2^)^2^*R*^2^. Besides the total stretching energy is *λ*(*δR/R*)^2^*R*^2^, where the strain on single cells scales as ∼ *δR/R*. These elastic deformations are a consequence of the external compressional pressure *P* that represents the constraining forces by the alginate capsules, and so the total work supplied to the elastic ring is ∼ *PR*^2^*δR*. The ring deformations are computed through the minimization of the total energy

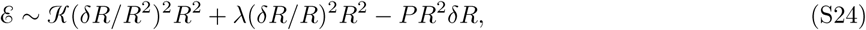

with respect to *δR*, yielding the compressive pressure as a function of the excess strain Δ*ϵ* = *δR/R*

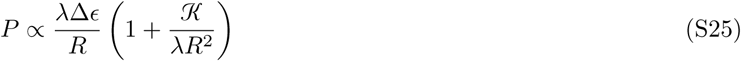

in terms of the geometrical and mechanical properties of the elastic ring.

For the parameter set in Table S1, the term *𝒦/λR*^2^ ≪ 1 is negligible, meaning that Eq. S25 is further simplified to *P* ∝ *λδR/R*^2^ for epithelial monolayers. Thereby, at the buckling transition for epithelia the excess strain Δ*ϵ* ∼ Δ*ϵ*_*c*_ takes the value given by Eq. S21, and thus the pressure at buckling reads

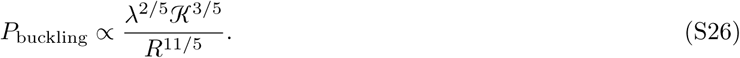

Similar arguments can be used to obtain the harmonic interaction between the elastic ring and the confinement in Eq. S2 and in particular the scaling of the spring constant *k* with the material properties of alginate capsules. Through similar arguments the reactive force of alginate capsules to an expanding internal pressure *P*\* can be computed, leading to an equivalent result to Eq. S25, except that the mechanics of the capsule are probed instead of the ring mechanics and the displacement *δR* is outwards. Consequently Eq. S25 turns into

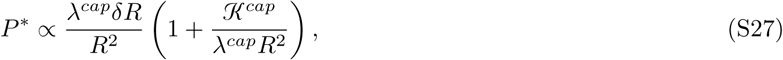

for a 3*d* spherical capsule confronted to an internal expanding pressure *P*\*. In consequence as we anticipated in Eq. S2, the capsule behavior resembles an harmonic potential with a spring constant *k* = *λ*^*cap*^*/R*^2^(1 + *𝒦*^*cap*^*/λ*^*cap*^*R*^2^), which according to the parameter set in Table S1 can be further approximated to *k* ∼ *λ*^*cap*^*/R*^2^.

#### Forces between confinement and an elastic ring after buckling

At the intermediate stages, the compressional forces accumulation results in a buckling transition, in which a macroscopic part of the tissue detaches from the inner surface of the capsule and folds. From this point on, the buckled tissue is not in physical contact with the capsule, except for a thin region at the edge through which compressive forces from the capsule may stabilize the tissue shape.

To further investigate the interplay between the buckled tissue and the compressional forces at the contact region, we assume that the tissue behaves as a 2*d* thin elastic ring with a bending modulus *𝒦* and compressional modulus *λ*. The following results has been reported previously elsewhere, (Chan and McMinn, 1966, pp. 433-442) in the main text. Here we revisit them in the context of tissue mechanics, but for more details about the intermediate calculations we refer to the original reference (Chan and McMinn, 1966, pp. 433-442) in the main text.

Let us consider a buckled elastic ring as illustrate in Fig. S6d, where the depth of the buckled ring is *δ* and the opening angle *α*. When adhesion is neglected, the part of the confinement in contact with the unbuckled segment of the ring withstands a uniform pressure *P*_*B*_ and a pair of normal forces per unit length *F*_*c*_ = *P*_*B*_*R* tan *α* at the contact points (red dots in Fig. S6d). These forces condition the shape of the inner elastic ring. In particular, enforcing force balance at the level of the contact points of the inner ring, results in a pair of compressive forces per unit length along the x-axis applied on the ends of the buckled ring, whose amplitude depends on the confinement-based forces through *F*_*B*_ = *P*_*B*_*R/* cos *α*. These external forces applied to the buckled ring cancel out, however, they are sufficient to buckle the ring and stabilize its shape. Therefore, the local force balance equation for a material element of the buckled ring reads

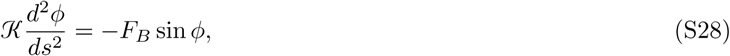

where *ϕ* is the angle between the tangent unit vector and the *x*-axis and *s* is the arc-length of the equilibrium shape. Thanks to the symmetry of the shape at the plane *x* = 0, the boundary conditions can be expressed as

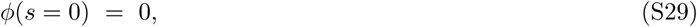

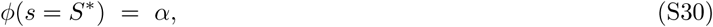

where 2*S*\* is the length of the buckled ring and 2*α* is the opening angle, Fig S6d. These set of equations determine a unique solution for *ϕ*, which is a complicated function of *α, S*\* and *F*_*B*_. However these physical quantities are not independent, but they are related to the excess strain of the elastic ring Δ*ϵ* = (*L –* 2*πR*)/2*πR* through conditions of self-consistency. In particular, at the contact points we enforce continuity of the curvature (i.e. *dϕ/ds* = 1*/R*) and continuity of the buckled ring (*x*(*s* = *S*\*), *y*(*s* = *S*\*)) = (sin *α*, cos *α*). Lastly, the excess strain is the sum of the axial strain of the unbuckled part and the difference in length of the buckled part compared with the original arc

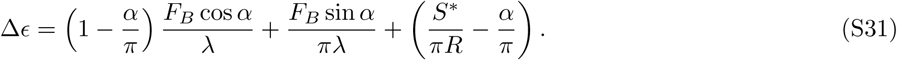

Notice that for *λ* → ∞, we recover the expected result for an inextensible ring.

For a given excess strain Δ*ϵ* and *λ*, there is a unique shape of the buckled ring that obeys all these conditions. These relations are transcendental and thus not analytically solvable. In Fig. S6g-i, we summarize some of the relevant geometrical properties of the buckled shape as a function Δ*ϵ* and *λR*^2^*/𝒦*: from left to right the depth of the buckled ring *δ*, the opening angle *α* and the dimensionless buckling forces *F*_*B*_*R*^2^*/𝒦*. All of them obey a power-law with the excess strain Δ*ϵ* and most remarkably to a good approximation they are independent of the compressional modulus *λ*. Consequently, the shape of the buckled ring provides a simple readout of the dimensionless mechanical forces at the contact line *F*_*B*_*R*^2^*/𝒦*.

Here, we detail the fitting method used to construct Figs. 4h-i in the main text. As explained above, by enforcing force balance, we obtain the triad of parameters (*α, δ/R, P*_*B*_*R*^3^*/𝒦* cos *α*) as a function of Δ*ϵ* and *λR*^2^*/𝒦* only. From these parameters, we computed the angle *γ* at which the curvature is zero defining an inflection point for the fold shape, as shown in Fig. 4d in the main text. In Fig. 4h in the main text, we plot the pairs (*α, δ*), and (*γ, δ*) for a single value of the dimensionless compressional rigidity *λR*^2^*/𝒦*, and compare it to experimental measurements without free parameters. In Fig. 4i in the main text, we plot the pairs (*δ/R, P*_*B*_*R*^3^*/𝒦*) for a single value of the dimensionless compressional rigidity *λR*^2^*/𝒦*, and from the comparison with the experimental measurements, we find the best fit at the bending rigidity *𝒦* ∼ 0.5 ± 0.2 *μN* · *μ*m.

#### Restoring force of indented elastic rings before buckling

In the following, we study the elastic forces of a system made of a spherical monolayer prior to folding that is adhered to the inner side of an spherical elastic capsule that is subjected to a normal force *F*_*I*_, mimicking the effect of an indenter. Let us approximate the previous system by two concentric 3*d* elastic shells. The inner (outer) shell of radius *R* describes the tissue (alginate capsule), whose mechanical properties are characterized by the bending rigidity *𝒦* (*𝒦*^*cap*^) and the compressional rigidity *λ* (*λ*^*cap*^).

The system deforms under the action of a small external point-like force *F*_*I*_, which is localized and directed normal to the surface, forming a small bulge with radius *d* = *Rα* and depth *δ*. This problem was addressed in (Landau and Lifshitz, 1975, Theory of elasticity) in the main text, and here, we restrict the analysis to *α, δ* ≪ 1. At the bulge, the curvature of both surfaces scales as *δ/d*^2^, and the total bending energy integrated over the deformed surface (∝ *d*^2^) is (∼*𝒦* + *𝒦*^*cap*^)(*δ/d*^2^)^2^*d*^2^. Besides, the strain at the bulge area scales as ∼ *δ/R* and the total stretching energy due to the external force is ∼ (*λ* + *λ*^*cap*^)(*δ/R*)^2^*d*^2^. Therefore, the work generated by the external load ∼ *Fδ* is thus transformed into elastic deformations near the region where the force is applied. The shape of the bulge is given by the condition that the total mechanical energy

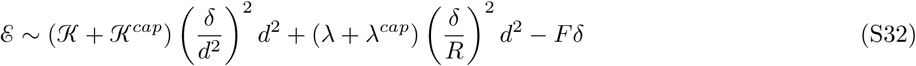

is minimal, yielding the following relation for the minimal radius *d* and depth *δ*

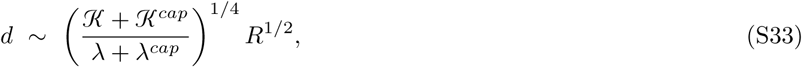

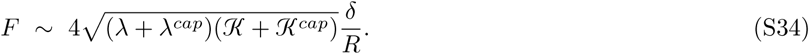

In conclusion, prior to the buckling transition the response of the system subjected to a point-like force is a restoring force proportional to the displacement of the bulged region *δ*. Hence, let us define two spring constant:

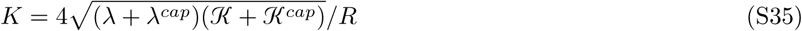

for capsules with unbuckled confluent monolayers and

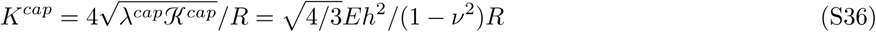

for capsules without tissues (*𝒦* = *λ* = 0).

To characterize the mechanical properties of epithelial tissues experimentally, we measure the linear regime of force-displacement curves by inducing small deformations with an indenter. Our findings permit to measure the spring constants under different configurations. In particular we compare the measured spring constant of capsules with and without tissues and we find that in general capsules are stiffer than the tissues underneath *𝒦* ≪ *𝒦*^*cap*^ and *λ* ≪ *λ*^*cap*^, meaning that the ratio between both coefficient can be approximated by

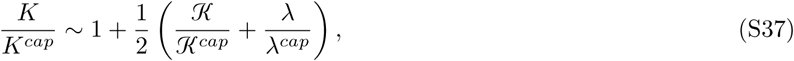

a simple relation that permits to decipher the mechanics of tissues.

### Section 5: Table of estimated parameters from the macroscopic approach

In this section, we summarize the main arguments used to determine the macroscopic parameters of epithelial monolayers that are bound to the inner surface of passive elastic shells in Table S1.

**TABLE S1:** Estimation of the material parameters of the physical model. Errors are SEM.

|  |  |  |
| --- | --- | --- |
| $E$ | Capsule elastic modulus | $19.5 \pm 0.7$ kPa (N=29) |
| $\nu$ | Capsule Poisson ratio | 1/2 |
| $h$ | Capsule thickness | $19.9 \pm 0.4$ $\mu\text{m}$ (N=54) |
| $R$ | Capsule radius | $97 \pm 1$ $\mu\text{m}$ (N=54) |
| $\mathcal{K}^{cap} = Eh^3/(12(1 - \nu^2))$ | Capsule bending rigidity | $17 \pm 1$ $\mu\text{N}\cdot\mu\text{m}$ |
| $\lambda^{cap} = Eh/(1 - \nu^2)$ | Capsule compressional rigidity | $0.52 \pm 0.02$ $\mu\text{N}/\mu\text{m}$ |
| $K^{cap}$ | Capsule without tissues spring constant | $0.064 \pm 0.003$ $\mu\text{N}/\mu\text{m}$ (N=30) |
| $K$ | Capsule with tissues spring constant | $0.074 \pm 0.007$ $\mu\text{N}/\mu\text{m}$ (N=16) |
| $\mathcal{K}$ | Tissue bending rigidity | $0.5 \pm 0.2$ $\mu\text{N}\cdot\mu\text{m}$ |
| $\lambda$ | Tissue compressional rigidity | $0.15 \pm 0.13$ $\mu\text{N}/\mu\text{m}$ |
| $\Delta\epsilon_c$ | Critical excess strain at the buckling transition | $\sim 0.1$ |
| $P_{\text{buckling}}$ | Pressure at the buckling transition | $\sim 100$ Pa |

The properties of alginate capsules are controlled externally. For the scale of tissue forces, alginate capsules behave as linear elastic materials with an elastic modulus *E* = 19.5 ± 0.7 kPa for 2.5% alginate capsules (Fig. 3 in the main text) and a Poisson ratio *ν* = 1/2. The typical thickness of the capsules is about *h* = 19.9 ± 0.4 *μ*m and their spherical shape has an inner radius of *R* = 97 ± 0.9 *μ*m. Since the thickness is smaller than the radius (*h* ≪ *R*), capsules are approximated by elastic shells with a bending rigidity *𝒦*^*cap*^ = *Eh*^3^/(12(1 – *ν*^2^)) = 17 ± 1 *μ*N · *μ*m and a compressional rigidity *λ*^*cap*^ = *Eh/*(1 *ν*^2^) = 0.52± 0.02 *μ*N*/μ*m, (Landau and Lifshitz, 1975, Theory of elasticity) in the main text. To estimate the macroscopic parameters of the monolayer, and in particular their bending and compressional rigidities (*𝒦* and *λ*), we require an additional independent measurement that is provided by the shape and mechanics of folded tissues. On the one hand according to Fig. S5g-i, there is a one-to-one *λ*-independent relation between the opening angle *α* of a buckled elastic ring and the dimensionless force at the contact points *F*_*B*_*R*^2^*/𝒦*, as both are invertible functions. The latter is related to the pressure on the capsule by *P*_*B*_ = *F*_*B*_ cos *α/R*. As explained in the main text by fitting the curve between the fold height *δ* and the capsule pressure *P*_*B*_, we obtain that the tissue bending rigidity is *𝒦* = 0.5 ± 0.2 *μ*N · *μ*m (see Fig. 4i in the main text), which is of the order of *𝒦* ∼ 10^8^ *k*_*B*_*T*.

When a point-like force is applied on a capsule, it forms a small bulge whose restoring force is different whether the inner surface is coated with a tissue or not. The slope of the force displacement curve defines an effective spring constant. The measured spring constant for capsules is *K*^*cap*^ ∼ 0.064 ± 0.003 *μ*N*/μ*m (see Fig. 4j in the main text). When the spring constant is measured for capsules with a confluent tissue, its value rises slightly to *K* ± 0.074 ± 0.007 *μ*N*/μ*m (see Fig. 4j in the main text), suggesting that the monolayer resist the applied load.

Next, we use the ratio between *K/K*^*cap*^ ∼ 1.16 ± 0.12 (see Fig. 4j in the main text) to obtain from Eq. S37 that *λ/λ*^*cap*^ ∼ 0.28 ± 0.24 as *𝒦/𝒦*^*cap*^ ∼ 0.03 ± 0.01 was determined above. Therefore, the compressional rigidity is *λ* = 0.15 ± 0.13 *μ*N*/μ*m for the value of *λ*^*cap*^ listed in Table S1.

A scaling argument allows to estimate the order of magnitude of the pressure at buckling *P*_buckling_. Let us consider that *𝒦* ∼ 1 *μ*N *μ*m, *λ* = 0.1 *μ*N*/μ*m and *R* ∼ 100 *μ*m, so that the dimensionless number *𝒦/λR*^2^ ∼ 10^−3^ ≪ 1. The threshold of the excess strain at which the circular elastic ring buckles is Δ*ϵ*_*c*_ ∼ 10^−1^, where the numerical factor has been set to 10 in Eq. S21. By introducing this value to the relation between the pressure and the excess strain S25, we obtain that the pressure at buckling *P*_buckling_ ∼ 100 Pa.

### Section 6: Description of a confined elastic ring (microscopic approach)

We showed in (Merzouki, Malaspinas and Chopard, 2016, pp. 4745-4754; Merzouki et al, 2018, pp. 511-519) in the main text how simulated growing cell monolayers can fold in the absence of external constraints. We found that the evolution of the cell monolayer morphology depends on the competition between the cell proliferation rate and the cell monolayer relaxation time. The cell mechanics affect directly the time needed by the cell monolayer to relax following a cell division, as well as the cell monolayer’s geometry, which are discriminant characteristics of folded tissues. In this work, we extend this buckling study by investigating the growth of cell monolayers inside elastic environments. Our simulations are compared to experiments, where spherical cell monolayers are cultured inside hydrogel microcapsules.

#### Cell monolayer cross-section

A circular cell monolayer cross-section is modelled by a ring of quadrilateral cells. Each cell has two lateral edges, which separate it from its neighbour cells, as well as two boundary edges, corresponding to the basal and apical edges. A cell monolayer is characterised by the area elasticity of its cells, the elasticity of its cell edges and the bending rigidity between two neighbour cells. The energy function *H* of the cell monolayer cross-section is the following:

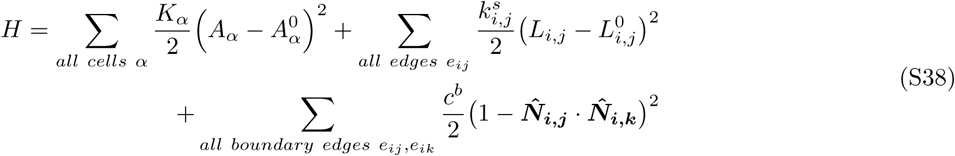

The first term of the energy represents the cell area elasticity. *K*_*α*_ and 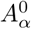 are the area elasticity coefficient and the preferred area of the cell *α*, respectively. In what follows, we set *K*_*α*_ = *K* = 10^9^ *N/m*^3^, (Merzouki, Malaspinas and Chopard, 2016, pp. 4745-4754; Merzouki et al, 2018, pp. 511-519) in the main text and 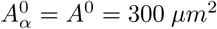 for all the cells *α*. The second term models the elasticity of the cell edges. 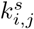 and 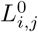 are the spring coefficient along an edge *e*_*ij*_ and its preferred (resting) length, respectively. The higher 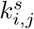, the more resistant are the cell edges to deformation (extension/compression). This term prevents cells from adopting triangular shapes under lateral compression. In what follows, we set 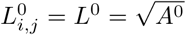 for all edges *e*_*ij*_. Finally, the third term stands for the bending rigidity of the cell monolayer, where *c*^*b*^ is the local bending rigidity coefficient. 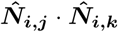 is the dot product between the vectors 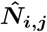 and 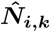. The latter are the unit vectors that are normal to the boundary (basal/apical) edges *e*_*ij*_ and *e*_*ik*_. This energy term is minimised when the normal vectors of neighbour cells are parallel and the tissue curvature is null.

By deriving this energy function *H* with respect to a vertex position ***r***_***i***_, we compute the internal force ***F***_***i***_ applied on the vertex *v*_*i*_.

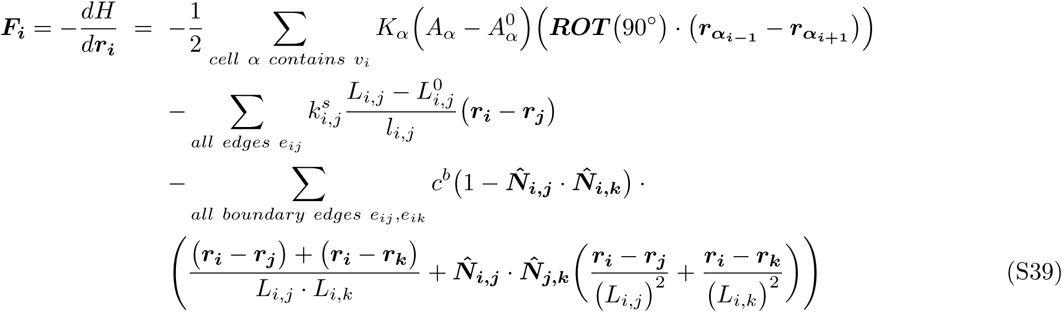

where

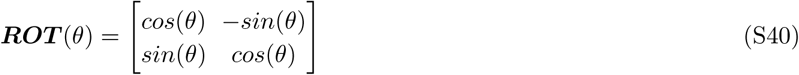

is the rotation matrix in two dimensions, and 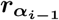 and 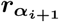 are the positions of the previous and next vertices, 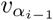 and 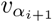, of the vertex *v*_*i*_ in the cell *α*, when the vertices of *α* are ordered counterclockwise

#### Heterogeneous elastic capsules

We constrain the cell monolayer cross-section using an elastic ring environment with a spring coefficient *k*_*ring*_.

This ring is characterised by its centre ***c***_***ring***_. The ring could have a homogeneous shape and its geometry be solely characterised by a radius *R*_*ring*_. However, for more realistic modelling of the experimental hydrogel micro-capsules, we actually constrain the growing cell monolayer cross-sections inside ‘circular’ elastic environments with heterogeneous borders. We perturb the circle radius with a set of sinusoidal signals. See Eq. (S41).

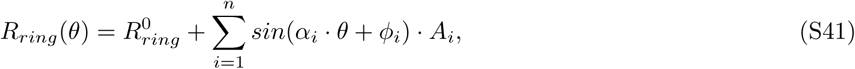

where *R*_*ring*_(*θ*) is the radius of the ring at an angle *θ, n* is the number of sinusoidal noises disturbing the initial radius of the ring 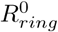, and *α*_*i*_, *ϕ*_*i*_ and *A*_*i*_ are the frequency, phase and amplitude of the *i*^*th*^ sinusoidal perturbation.

The contact of the cells with the inner surface of the capsule yields a friction that restrains the movement of the cells. Cells are subjected to non-slipping forces along the capsule border. In our model, when a vertex *v*_*i*_ at a position ***r***_***i***_ is close to the border of the capsule, ∥***r***_***i***_ − ***c***_***ring***_ ∥*> R*_*ring*_(*θ*_*i*_) – *ϵ*, it is subjected to a force that resists its tangential force component 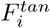 up to a given force threshold *F*_*NS*_, where the subscript *NS* stands for “Non-Slipping”. In contrast, when the vertex is far from the capsule border or when its tangential force component 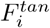 exceeds the non-slipping force threshold *F*_*NS*_, then the vertex is free from the non-slipping force exerted along the capsule’s surface.

### Section 7: Folding quantification

For each simulation, the contour of the confining capsule is stored as well as the basal contour of the cell monolayer over time. The contours are stored in .csv files. Each line contains the (*X, Y*) coordinates of a point along the contour. Points are ordered counter-clockwise. The basal contour of the cell monolayer is defined by the counter-clockwise sequence of basal vertices *v*_*i*_, at positions ***r***_***i***_ = (*x*_*i*_, *y*_*i*_), along the cell monolayer cross-section. The dynamics of the cell monolayer can then be followed over time.

From the sequence of vertex positions ***r***_***i***_ = (*x*_*i*_, *y*_*i*_), we compute the radius *R*_*i*_ = ∥***r***_***i***_−***c***_***ring***_∥ of each vertex (distance to the capsule center ***c***_***ring***_) and its angle *θ*_*i*_ along the cell monolayer contour. The first characteristic property of a fold is its detachment from its initial unfolded configuration and its displacement from the border to the interior of the capsule. In order to identify a detachment, we compute the difference between a given contour profile *R*(*θ*) and the reference profile *R*_*ref*_ (*θ*); *dist*(*θ*) = *R*(*θ*) – *R*_*ref*_ (*θ*). The reference profile corresponds to the initial unfolded profile subjected to the confinement constraints. When the tissue detaches from the capsule and moves inward, the distance *dist*(*θ*) to the reference profile in this region is negative; *dist*(*θ*) *<* 0. In contrast, when *dist*(*θ*) *>* 0, this corresponds to a tissue that pushes outward on its confining environment. Because the number of cells (and vertices) of the growing simulated tissue evolves over time, the comparison between the tissue profiles requires a pre-processing. All the profiles are first interpolated to get their radius at specific angles *θ* ∈]0, 2*π*].

The identification of detachments along the tissue contour starts with finding the local minima of the distance function with respect to the angle *dist* = *f* (*θ*). In order to ignore noises and minor detachments, we set a threshold *ϵ*_*dist*_, ex. *ϵ*_*dist*_ = –10*μm*, above which a local minimum is dismissed. The second step aims at delimiting the detachment, i.e. finding its start and end points. From each significant distance local minimum, we look backward and forward along the tissue contour for points with *dist* ≈ 0, ex. *dist > –* 1*μm*. These start and end points allow us to compute the width of the detachment. Moreover, the position of a detachment is computed as the mean position between the start and end points delimiting the detachment. In case different distance local minima have the same start and end points, they are considered as potential folds being part of a unique detachment.

There, we distinguish between a detachment and a fold. The fold delimitation is not solely based on the distance to the reference contour, but it is also characterised by an increased local curvature. The local curvature is the second derivative of the distance function *dist* = *f* (*θ*). In summary, a fold is characterised by a distance to the capsule and a curvature which exceed given thresholds *ϵ*_*dist*_ and *ϵ*_*curv*_, respectively. In what follows, we use *ϵ*_*dist*_ = –10*μm* and *ϵ*_*curv*_ = 5*e*^−4^. The start and end points of a fold are the points along the contour (before and after the fold position) where the first and second derivatives of the distance change sign.

### Section 8: Simulations and results

#### Effect of cell mechanics on buckling

In this section, we show how different cell and tissue mechanical properties, namely the cell stiffness and the tissue bending rigidity, affect the timing of tissue folding as well as the number, the width and the position of folds that form.

For this purpose, we simulate 45 cell monolayer cross-sections for a set of normalised parameters 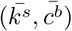, where 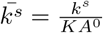 and 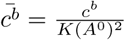. For each couple of normalised parameters 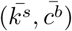, we start with a relaxed ring of 40 cells. All the cells are assigned the same preferred area *A*^0^ = 300 *μm*^2^ and an area elasticity coefficient *K* = 10^9^ *N/m*^3^. The cell monolayer cross-section is confined inside an elastic heterogeneous ring, with *k*_*ring*_ = 0.06 *N/m* and a non-slipping force 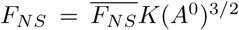 with 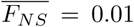, which corresponds to *F*_*NS*_ = *O*(10^−8^) *N*. The radius of the heterogeneous ring is perturbed using *n* = 5 sinusoids with frequencies *α*_*i*_ = *i*, ∀*i* = 1..*n*, random amplitudes *A*_*i*_ ∈] 0*…*0.02 *R*] and random phases *ϕ*_*i*_ ∈] 0*…*2*π*]. Fig. S7a presents the distribution of the number of folds forming after 40 cell divisions and the number of cells when the first fold appears for each set of normalised parameters 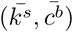.

We observe that higher bending rigidity 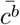, which increases the differential tension between the boundary and the lateral cell edges, enhances cell monolayer thickening and therefore postpones and reduces tissue folding. In contrast, higher cell stiffness 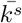, which resists cell compression, accelerates and enhances tissue folding in response to cell proliferation.

In Fig. S7a, we highlight in red the histograms matching the experimental observations, i.e. a majority of 2 folds formation, followed by cases where 1 fold is observed, then fewer cases displaying 3 folds. This parameter region corresponds to a first fold forming at approximately 70 cells and an average cell aspect ratio at buckling *AR* = 2 – 2.5, while regions with larger bending rigidity show a postponed buckling when cells are much more elongated (*AR* = 3 – 6). We found that along the line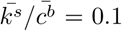, the fold number (Fig. S7a) was similar to experiments (see Fig. 5 in the main text).

The forces applied by the tissue on the capsule (and the other way around, the external forces applied by the capsule on the tissue) over time were monitored during our simulations. The evolution of the estimated pressure over time are presented in Fig. S6b. Experimentally, the force applied by the growing tissue on the capsule at buckling is estimated to be in the order of the micronewton *O*(*μN*) (*F* = 0.4 – 4 *μN*), and the pressure to be in the order *O*(0.1 *kPa*). In the highlighted regions of the model parameters (matching the experimental distribution of the number of forming folds), the first folds appear in our simulated tissues when the number of cells reaches ≈ 60 −70. In this time interval, the force applied by the simulated tissue cross-section on its elastic environment 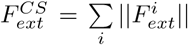 varies between 0.5 *μN –* 1 *μN*. The force 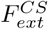 is the sum of all the external force norms applied by the tissue cross-section on the capsule. In 3D, we estimate the force applied by the spherical tissue 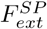 on the capsule to be:

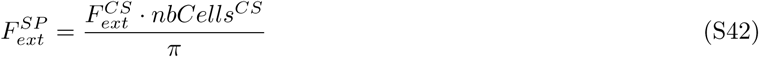

Therefore, the pressure *P* at buckling in our simulations is computed as 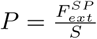, where *S* = 4*πR*^2^ is the surface of the sphere with radius *R*. In our simulations, *R* ≈ 100*μm*. Experimentally, the capsule radius varies between 75*μm* and 175*μm*, which corresponds to a 150*μm –* 350*μm* diameter. Finally, with approximately 70 cells at buckling, and a force applied by the tissue cross-section 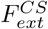 between 0.5 *μN* and 1 *μN*, we estimate that the pressure at buckling in our simulations to be *P* ≈ 0.17 *kPa* for values of *k*^*s*^ ∼ 3 10^−3^ *μ*N/*μ*m and *c*^*b*^ ∼ 9 *μ*N *μ*m Fig. S6b. This is consistent with the experimental pressure in the order of 0.1 *kPa* estimated when the cultured cell monolayers buckle inside the elastic microcapsule.

#### Effect of confining environment

The non-slipping forces applied by the capsule on the cells play an important role on the timing, the number and the shape of the folds that form. We analysed the folding of simulated cell monolayers growing inside capsules with normalised non-slipping forces 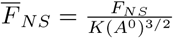 varying between 0.001 and 0.01.

We found that an increased friction between the capsule’s inner surface and the cell monolayer leads to accelerated folds formation. More folds appear and they present wider shapes. In general, once a fold forms, the lateral cell compression in this region decreases. Without friction, the cells around the fold migrate into the fold, leading to the constriction of the fold’s neck and the formation of a highly curved inward bud. In contrast, increased non-slipping forces prevent the migration of the cells into the fold that just formed. Therefore, the fold grows wider, other parts of the cell monolayer keep on accumulating lateral compressive stress due to cell proliferation, and additional folds form. Click here to watch the video, or follow the link: https://youtu.be/C73R6kLNfQk.

## Captions for Movies 1-6

**Movie 1: MDCK H2B-eGFP mCherry-Actin monolayers forming one fold (left panel) and two folds (right panel) inside a** 2.5% **alginate capsules.** Red channel is mCherry-Actin, green channel is H2B-eGFP and cyan is Alginate is labelled with ATTO647. Time-lapse is 2.5 hours. Scale bar, 100 *μ*m

**Movie 2: A cell monolayer treated with 10** *μ***M Blebbistatin.** Red channel is mCherry-Actin, green channel is H2B-eGFP and cyan is Alginate is labelled with ATTO647. Time-lapse is 2.5 hours. Scale bar, 100 *μ*m.

**Movie 3: A cell monolayer treated with 10** *μ***M Mitomyocin C.** Red channel is mCherry-Actin, green channel is H2B-eGFP and cyan is Alginate is labelled with ATTO647. Time-lapse is 2.5 hours. Scale bar, 100 *μ*m.

**Movie 4: Cell monolayer relaxation over short period of time after capsule dissolution with alginate lyase corresponding to the top panel of Fig. 2c.** Red channel is mCherry-Actin, green channel is H2B-eGFP and cyan is Alginate is labelled with ATTO647. Time-lapse is 15 seconds. Scale bar, 100 *μ*m.

**Movie 5: Deformation of a** 1% **alginate capsule over time resulting from MDCK H2B-eGFP mCherry-Actin monolayer buckling.** Red channel is mCherry-Actin, green channel is H2B-eGFP and cyan is Alginate is labelled with ATTO647. Time-lapse is 2.5 hours. Scale bar, 100 *μ*m.

**Movie 6: Simulated cell monolayer forming one, two, three, four and five folds (from left to right, respectively) inside an elastic capsule (***k*_*spring*_ = 0.06 *N/m*). The simulation were performed within the model parameters region matching the experimental fold number distribution. The normalised non-slipping force between the growing cell monolayer and the underlying capsule is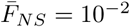, which corresponds to *F*_*NS*_ = *O*(10^−8^ *N*) with *K* = 10^9^ N/m^3^ and *A*^0^ = 300 *μ*m^2^. The colours represent the cell’s aspect ratio *AR* = width/height. Red corresponds to high *AR* ≈ 1, where width ≈ height. Blue corresponds to elongated cells with low *AR*, where width ≪ height.

